# Molecular sources of monoterpenoid chemodiversity in the Asteraceae *Tanacetum vulgare* suggest a new model for the evolution of specialized metabolism

**DOI:** 10.64898/2026.08.18.745266

**Authors:** Marvin Hildebrandt, Bianca Laker, Dominik Ziaja, Elisabeth Eilers, Prisca Viehöver, Ruth Jakobs, Stephan Hammer, Tobias Busche, Marion Eisenhut, Caroline Müller, Andrea Bräutigam

## Abstract

Highly diversified specialized metabolism enables plant communication with pollinators, herbivores, and protectors^1–4^. Its chemodiversity, under which evenness, richness and variation is summarized^5–7^, includes many compounds without known function^1–4^ and presents an evolutionary conundrum about how and what is selected for^8,9^. Due to its complexity, it is frequently unknown how it is encoded in genomes. To produce population level chemodiversity, the traits need to allow for highly chemodiverse and highly specific individuals in the same population. Here we use metabolomics, transcriptomics, and genomics combined with field analyses and functional assays of monoterpene synthases in the Asteraceae *Tanacetum vulgare* (tansy) and identify forces which produce high population level chemodiversity: selection for product specificity in enzymes, loss-of-expression alleles, absence variation, and specialized metabolism islands drive individuals towards low chemodiversity while unlinked enzyme loci, expression variation alleles, presence variation, and *de novo* enzyme evolution enable high individual chemodiversity. Since the molecular data suggests selection for mechanisms that increase chemodiversity itself at the population level, the screening hypothesis which posited plants produce a reservoir of diverse chemicals prior to selection^8^ should be replaced by a chemodiversity selection hypothesis. The results demonstrate that, in addition to plant protection via individual chemicals with known targeting mechanisms for predators, being different from your neighbors even if you are closely related is likely an important element in plant protection.

## Introduction

Plants produce a broad range of specialized metabolites, which play an important role in interactions with both antagonistic and beneficial organisms^1–4^ and generally enhance various reproduction-related traits^2–4^. The specific composition of specialized metabolites generated by the genome architecture in each organism influences its survival and fitness. Diverse ecosystems with a patchy distribution of specialized metabolites across plant individuals generally better withstand pathogen and herbivore pressure compared to agricultural monocultures^1^. Verbal, quantitative and empirical models including the famous screening hypothesis which posits plants produce a reservoir of diverse chemicals prior to selection have been put forward to explain the evolution of chemodiversity within a species^7–9^. We hypothesized that a population level analysis reveals the molecular trait architecture of such chemodiversity^10–12^ and provides support for a new model.

*Tanacetum vulgare* (tansy, Asteraceae) is an invasive weed^13,14^ to North America and a model plant for population level and individual chemodiversity^14^. Chemodiversity in leaves and flowerheads of the obligate outcrosser *T. vulgare* is largely driven by qualitative and quantitative variation in monoterpenoids, which shapes interactions with herbivores, their natural enemies, and pollinators^2,4,15,16^. Consequently, intraspecific differences in monoterpenoid profiles have been shown to influence plant performance, community assembly and possibly the invasion success^2,3,14–16^. Monoterpenoid compositions in *T. vulgare* leaves are stable across clones and can significantly differ between individuals even under controlled environmental conditions, which lead to the definition of “chemotypes”^17,18^. The structure of monoterpenoids is based on a ten-carbon backbone synthesized by a prenyltransferase from isopentenylpyrophoshate and dimethylallylpyrophosphate which are condensed to geranylpyrophosphate (GPP). Monoterpene synthases (MTSs) with different plasticity regions in their protein sequence convert GPP to a diverse array of linear, cyclic, and bicyclic monoterpenes^19^. Evolution of substrate specificity is frequently convergent so that products cannot be inferred from phylogeny. Crossing experiments with the obligate outcrosser *T. vulgare* have revealed partially non-Mendelian inheritance of monoterpenoid profiles^17^. Analyses in other species, general modelling approaches^9^, and meta-analyses suggest multiple modes that may enable high chemodiversity^8^.

To determine the molecular mechanisms which can generate both highly diverse and comparatively specific monoterpenoid individual chemotypes within a single, local population of *T. vulgare*^2^, we combined field sampling of the local population, population-wide metabolomics and transcriptomics, functional enzyme assays, genome sequencing, and targeted enzyme mutagenesis. High intraspecific chemodiversity in the local population and the detection of molecular mechanisms for both increasing and decreasing individual chemodiversity suggest that variation potential is built into the trait and therefore chemodiversity itself is selected for. Understanding the molecular trait architecture of chemodiversity provides the tools for future manipulation of population-level chemodiversity in crops to improve protection.

## Results

To establish a minimal set of likely necessary MTS enzymes responsible for the chemodiversity in tansy, a local population was characterized in detail. In leaves of 39 individuals field-sampled within an area of ca. 1 km^2^ near Bielefeld, Germany, at least 32 monoterpenoids were detected in total, with variable composition in each individual, and ranging from complete absence to contributing up to 90.2% of the total blend (Figure 1A, Extended data Fig. S1). Seven hydrocarbons (santolina triene, α-pinene, β-pinene, camphene, sabinene, γ-terpinene and p-cymene), eight alcohols (borneol, verbenol, *cis*-chrysanthenol, *trans*-sabinol, *trans*-sabinene hydrate, *cis*-sabinene hydrate, artemisia alcohol and yomogi alcohol), eight ketones (α-thujone, β-thujone, chrysanthenone, camphor, pinocarvone, piperitone, umbellulone and artemisia ketone), seven esters (β-carvylacetate, *trans*-sabinyl acetate, α-chrysanthenyl acetate, *cis*-verbenol acetate, *trans*-β-chrysanthenyl acetate, terpinen- 4-yl acetate and β-artemisia acetate) and two ethers (1,8-cineole and myroxide) were detected (Figure 1A, Extended Data Fig. S1).

**Fig. 1:**
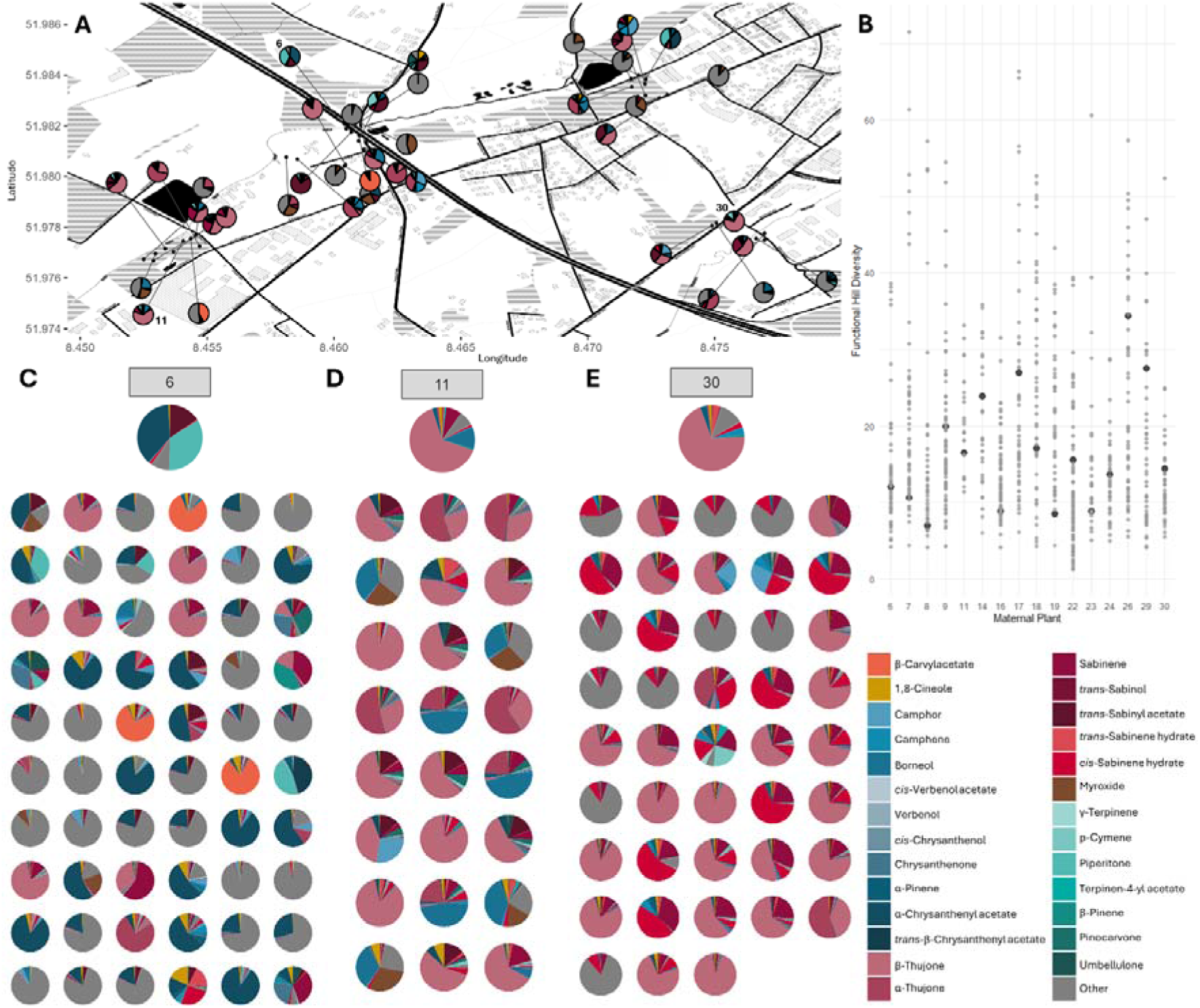
Characterization of leaf monoterpenoid chemodiversity from plants of a local *T. vulgare* population (maternal plants) and their descendants. A, Sampling area with locations and leaf monoterpenoid profiles of sampled plants. **B**, Functional Hill diversity of maternal parents (black) and their descendants (grey). **C**, Leaf monoterpenoid profiles of (maternal) plants no. 6, 11, 30 and their descendants. All other profiles provided in Figure S2.

While the monoterpenoid profiles clustered into seven chemotypes based on dominant constituents (Extended data Fig. S3), the distribution and relative abundance of monoterpenoids was continuous. The functional Hill diversity of the monoterpenoids in these 39 plants which compresses chemodiversity into a single number based on metabolite abundances and similarities ranged from 4.3 to 38.4 (Figure 1B). To establish a *T. vulgare* collection grown in controlled conditions, seeds were collected from the same plants, 849 descendants were grown in a greenhouse, and leaf monoterpenoid profiles were measured (Figure 1C-E and Extended Data Fig. S2). Maternal plants and descendants were both numbered and designated by their origin; 11-7 for example refers to the 7^th^ descendant of maternal plant 11. Descendants were highly diverse compared to each other and their maternal parent (Figure 1C-E). The two major monoterpenoids piperitone and α- chrysanthenyl acetate in maternal plant 6 were retained in only three descendants and β-carvylacetate was the dominant monoterpenoid in 3 out of 60 descendants, although this compound was absent from maternal plant 6 itself (Figure 1C). Stable transmission of the chemotype to some of the offspring was observed in some cases, with 11 of 24 descendants of maternal plant 11 being also dominated (> 50%) by β-thujone (Figure 1D). *Cis*-sabinene hydrate was the dominant monoterpenoid (> 50%) in 6 of 43 descendants of maternal plant 30 that itself contained only 2.8% *cis*-sabinene hydrate (Figure 1E). Functional Hill diversity of descendant plants varied between 1.2 and 71.5, with highly diverse maternal plants being able to yield comparatively low diversity descendants and vice versa (Fig. 1B). Together with the observation of 19 low diversity individuals dominated by a single compound (> 50%), the variation between parents and descendants suggests that mechanisms exist which enable both high and low diversity individuals in the same population.

To investigate these molecular mechanisms, twenty-four descendant lines from 13 different maternal plants with functional Hill diversities from 6.4 to 45.2 representing all 32 monoterpenoids initially detected were clonally propagated for further analyses. The biggest contribution to the functional Hill diversity of monoterpenoids in those plants resulted from differences in evenness but not disparity or richness pointing to quantitative differences (Extended Data Fig. S5). Structural analyses suggested that of 32 identified monoterpenoids five (santolina triene, artemisia alcohol, yomogi alcohol, artemisia ketone and β-artemisia acetate) were likely products of prenyltransferases or products of MTS acting on substrates other than GPP^20,21^ (Figure 2A, Extended Data Fig. S6). The remaining 27 monoterpenoids were likely generated from 13 monoterpenes (β-ocimene, limonene, 1,8-cineole, γ-terpinen, terpinen-4-ol, camphene, bornyl diphosphate, α-pinene, β-pinene, sabinene, *cis*-sabinene hydrate, *trans*-sabinene hydrate and α-thujene) by oxyfunctionalization, and/or oxidoreduction, and/or acetylation (Figure 2A, Extended Data Fig. S6). At least six of the MTS are product-specific (α-pinene, sabinene, limonene, camphene/bornyl diphosphate, *cis*- sabinene hydrate, β-ocimene producing MTS), since lines that nearly exclusively contain monoterpenoids derived from that MTS product are detected (Figure 2A). Hence product-specific enzymes are selected for in *T. vulgare*.

**Fig. 2:**
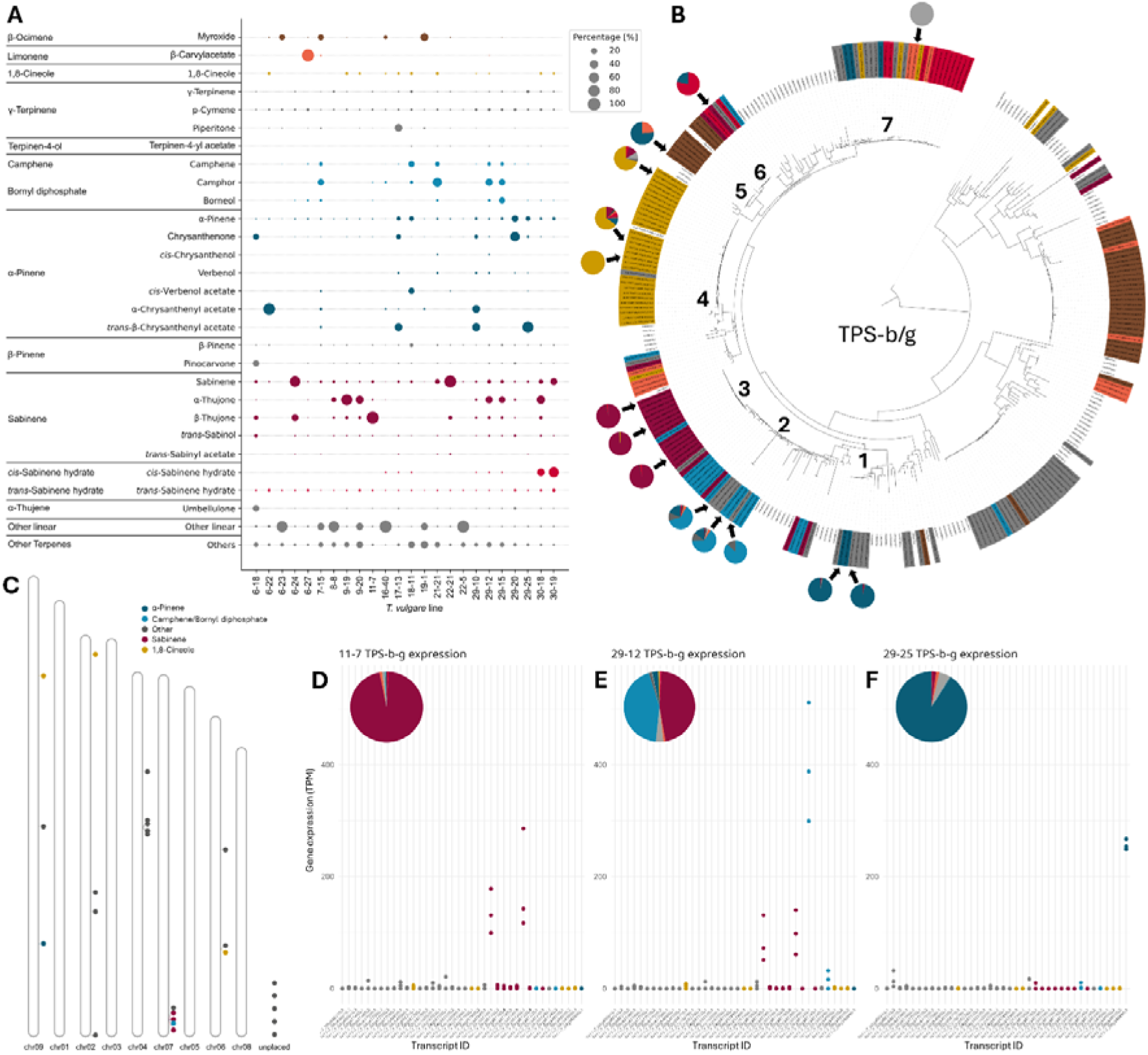
Monoterpenoid profiles of 24 *T. vulgare* chemotypes can in part be explained by expression divergence of seven MTS types. **A**, Monoterpenoid profiles of leaves of 24 propagated lines sorted by MTS product. **B**, Phylogenetic tree of MTS candidate transcripts and product profiles identified by in vitro enzyme assays. Branches with validated MTSs are numbered. Color coding analogous to Fig. 1. **C**, Locations of MTSs in the genome of line 11-7. **D-F**, Expression of MTSs identified in the 11-7 genome for three clones per line and leaf terpenoid profile for lines 11-7, 29-12 and 29-25. Leaf terpenoids derived from the same MTS product are summed.

To identify the molecular identity of the MTSs underlying the observed chemodiversity, transcriptomes of developing leaves of the 24 propagated lines (Figure 2A) were sequenced to a depth of ca. 50 million reads per line and transcriptomes were assembled. 1,699 *T. vulgare* transcripts from 684 genes contained Pfam terpene synthase (TPS) domains PF01397 and PF03936 (Extended Data Fig. S7 and Supplementary Table 1). A phylogeny of *T. vulgare* TPS candidates with TPSs from the Asteraceae *Cynara cardunculus*, *Helianthus annuus*, and *Artemisia annua*, with TPSs from *Arabidopsis thaliana*, and with TPSs from *Marchantia polymorpha* (Extended Data Fig. S7) identified 201 candidate *T. vulgare* (Tv) monoterpene synthases (MTSs) of the phylogenetic groups TPS-b and TPS-g which typically contain MTSs (summarized in Jiang et. al. 2019^22^) (Figure 2B). Seven branches of this phylogeny were analyzed in detail. In branch 1, seven TvMTS candidates derived from six lines clustered with three *A. annua* (Aa) MTS candidates (bootstrap support 100%) and Spearman correlation of transcript abundances with monoterpenoid abundances suggested α-pinene or a monoterpene for which no specific MTS is expected („other”) as the likely product (Figure 2B). Functional analyses of two enzymes yielded >96% α-pinene as the product. 17 TvMTS candidates in branch 2 (bootstrap support 19%) and 15 TvMTS candidates in branch 3 (bootstrap support 100%) clustered with a camphene synthase and eight additional MTS candidates from *A. annua* (bootstrap support 95%, Figure 2B and Extended Data Fig. 7). Correlation analyses with monoterpenoid abundances suggested camphene and bornyl diphosphate as the likely products of TvMTS candidates in branch 2 and sabinene as the likely product of TvMTS candidates in branch 3 (Figure 2B). Functional analyses of three enzymes from each branch yielded >95% sabinene as the product of TvMTS candidates in branch 3 and 62-87% camphene with substantial production of α-pinene in two out of three cases for TvMTS candidates in branch 2 (Figure 2B). Of the 32 TvMTS candidates in branch 4 that correlated best with 1,8 cineole and co-clustered with two AaMTS candidates (bootstrap support 99%), three were functionally tested and confirmed as 1,8-cineole synthases whose specificity ranged from 65% to 100% (Figure 2B). Six MTS candidates in branch 5 (bootstrap support 100%) clustered with no AaMTS candidates and one tested candidate yielded 76% α-pinene and 24% limonene making this a phylogenetically unrelated second α-pinene synthase in *T. vulgare* (Figure 2B). Six MTS candidates in branch 6 (bootstrap support 100) clustered with one AaMTS candidate and one tested candidate yielded 78% *cis*-sabinene hydrate and 22% α- pinene (Figure 2B). One candidate from branch 7 yielded the non-linear monoterpene linalool (Figure 2B). The molecular identity of MTSs producing sabinene, camphene, α-pinene, 1,8- cineole, limonene, *cis*-sabinene hydrate, and linalool was thus established via functional *in vitro* assays. Closely related enzymes generally produce the same main product and can be both highly specific and broad spectrum enzymes contributing to low and high diversity, respectively. The isolation of transcriptome-based candidate MTSs (Figure 2B) which are not or barely present in the monoterpenoid chemotype profiles (Figure 2A) suggested that low transcript abundance of MTS also contributes to low diversity.

To test the potential contribution of presence absence variation to terpenoid chemodiversity, the number of MTS loci in the genome of a low chemodiversity line, the genome of line 11-7 that is specific to the sabinene-derived terpenoid β-thujone (90% of terpenoid content), were analyzed by genome sequencing. 10,257,307 long reads (182.15 Gbp) were assembled into 58,151 contigs with an N50 of 299.24 kbp and a total length of 6.81 Gbp, separated into two haplotype assemblies and scaffolded into nine chromosomes for each haplotype assembly using the Darwin Tree of Life release of a UK *T. vulgare* individual with unknown chemotype^23^ (Figure 2C). 20 MTS candidates were placed on five of the nine chromosomes and five MTS candidates remained on unplaced contigs, making them candidates for presence absence variation. The camphene synthase from branch 2 and three sabinene synthases from branch 3 (Figure 2B) were detected within 1.5 Mbp on chromosome 7 (Figure 2C) and possibly result from duplication events. Back mapping of the transcriptome data of three clones of line 11-7 to its genome demonstrated high expression of two out of three genes (Figure 2D). The very low-level transcript abundance <21 TPM (transcripts per million) of all other MTS enzymes explained why they were assembled from the transcriptome (Figure 2B) without leading to detectable product formation (Figure 2A). Transcriptomes from three clones of line 29-12 which contains sabinene- (40%) and camphene-/bornyl-diphosphate-derived (38%) monoterpenoids show high transcript abundance of two sabinene synthase transcripts and one camphene synthase transcript (Figure 2E). In line 29-25, which is dominated by α-pinene-derived monoterpenoids (87%), high transcript abundance of the α- pinene synthase transcript is detected in all three replicates (Figure 2F). Flux through MTS from the common substrate GPP is hence likely controlled by enzyme abundance derived from transcript abundance. The results support that expression differences are indeed a major factor for determination of chemotype complexity in the individual and therefore chemodiversity in the population.

To test the contribution of neofunctionalization to population chemodiversity, sabinene and α-pinene production were investigated in detail. The phylogeny suggested that the sabinene synthase evolved from a progenitor camphene synthase (Figure 3A), given that the gene in the last common ancestor of *A. annua* and *T. vulgare* from which the tree branch originated likely was a camphene synthase^24^ (Figure 3A). A synteny analysis of the MTS-containing region of *A. annua* (Aa) contig PKPP01000369 and *T. vulgare* (Tv) chromosome 7 identifed the regions as synthenic since the MTS candidate genes in *T. vulgare* are surrounded by the genes PRP19, TRM32, two genes of unknown function, ARPC2B, CYN and U2AF2 which also surrounded the MTSs in *A. annua* (Figure 3B). In *T. vulgare* this genome region also contained two candidate oxyfunctionalization genes, a P450 and a peroxidase, and five candidate dehydrogenases. The region in *A. annua* only contained one candidate dehydrogenase and two caleosin-related proteins, which might also be involved in oxyfunctionalization. This suggested that the region is a conserved specialized metabolism island, which expanded in *T. vulgare* and provided a basis from which the highly specific sabinene synthase evolved.

**Fig. 3:**
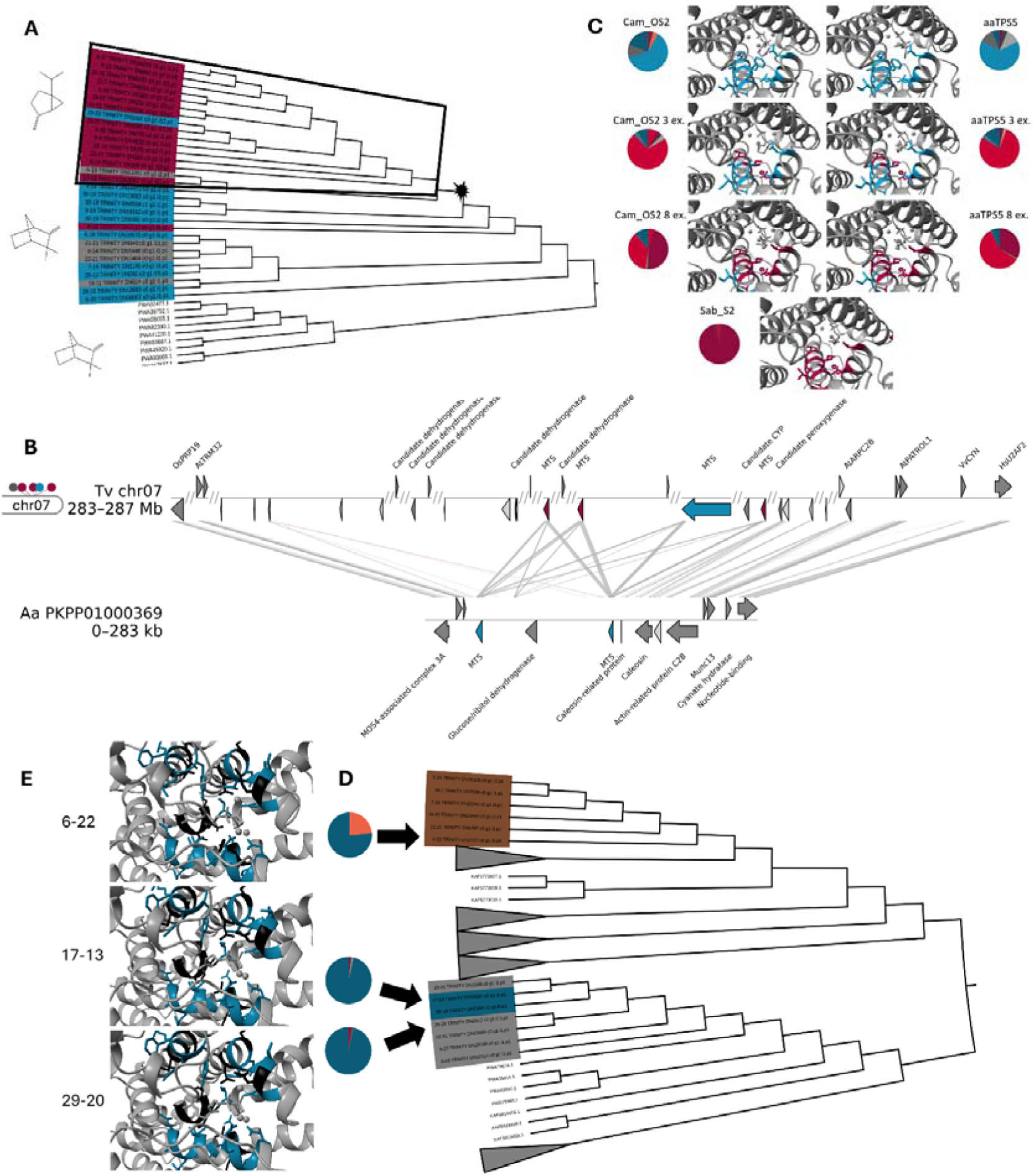
Chemodiversity in T. vulgare evolved through at least two neofunctionalizations requiring non-canonical mutations. **A**, Detail of figure 2B containing functional camphene and sabinene synthases. **B**, Genomic region with camphene/sabinene synthase tandem duplication in *T. vulgare* line 11-7 compared to the corresponding region in *A. annua*. Colored arrows depict matches to characterized MTSs and dark grey arrows depict conserved syntelogous genes between both regions. All other annotated genes are shown in light grey. Regions without genes were omitted (marked by “//”) **C**, Binding pocket and *in vitro* product profile of WT camphene and sabinene synthases and mutants with three and eight exchanged amino acids show changes in binding pocket geometry alters product profiles. **D**, Branches 1 and 5 of the phylogenetic tree showing transcripts and in vitro product profiles of individually evolved α-pinene synthases. **E**, Binding pocket of individually evolved α- pinene synthases.

Protein alignments of the camphene and the sabinene synthases from the *T. vulgare* transcripts in branches 2 and 3 detected 20 amino acid changes between the two enzyme groups of which three reside in known plasticity/binding sites known to be relevant for the function (amino acid positions 321, 345 and 348^19^). Five changes lay in helices adjacent to the active site (Extended Data Fig. S8). The camphene synthase of *T. vulgare* and the camphene synthase of *A. annua* (Figure 3A) produced comparable product profiles dominated by camphene and shared all of the 20 diagnostic changes detected between *T. vulgare* enzymes (Figure 3C, Extended Data Figure S8). Mutation of the three residues in the plasticity regions^19^ shifted the product profile in an *in vitro* enzyme assay to sabinene hydrate instead of sabinene in both the *T. vulgare* and the *A. annua* enzymes (Figure 3C). Additional mutation of the five residues in helices adjacent to the binding pocket resulted in 32% and 46% sabinene production, respectively. The camphene synthases thus have the potential to evolve via a short path into a sabinene synthase and increase chemodiversity in a species. The wild-type sabinene synthase from *T. vulgare* achieved 95% sabinene production (Figure 3C) which indicated that while active site and active site adjacent mutations are sufficient to access sabinene production, selective pressure has optimized the enzyme to near specificity by also modifying the shell region.

Genome analyses (Figure 2C) placed only one of two α-pinene synthases (Figure 2B) on the 11-7 genome making the second α-pinene synthase detected from 6-22 another candidate for presence absence variation. A detailed phylogeny confirmed these enzymes on two distant branches of the phylogenetic tree, branch 1 and branch 5 (bootstrap support 100%, Figure 3D). Branch 1 enzymes co-localized with the β-pinene synthase QH6 from *A. annua.* Branch 5 enzymes did not cluster together with candidates from the other Asteraceae species tested but co-localized with branch 6 that contained the uncharacterized TPS7 from *A. annua* (UniProt ID A0A165D1P5) and a sabinene hydrate synthase from *T. vulgare* (Figure 2B). Template modeling and comparison of both clusters of α-pinene synthases showed that the active sites contain 18 identical and 15 unconserved amino acids in the NSE/DTE motif and plasticity regions (Figure 3E, Extended Data Figure S12) and suggested that a suite of different amino acids may confer a geometry of the active site which enabled α-pinene production. *T. vulgare* thus contains at least two MTS neofunctionalizations since the recent divergence of *A. annua* and *T. vulgare* (4.5-12.7 million years ago^25^).

## Discussion

Since specialized metabolism evolves rapidly^26,27^, chemodiversity at the population level represents a mini-laboratory of evolution. To create population level chemodiversity, the population needs to contain plants with high and with low individual chemodiversity. The molecular architecture provided evidence for mechanisms increasing and decreasing *individual* chemodiversity at the protein, the transcript, and the genome level.

*T. vulgare* has evolved a highly specific sabinene synthase from an enzyme in the last common ancestor which produced camphene (Figure 2). Phylogeny, functional testing (Figure 2B) and mutational analyses (Figure 3A, B) charted the evolutionary path. In addition to amino acids near the enzyme core^19^ (Figure 3B), changes in the shell were required for evolving a sabinene synthase with high specificity (Figure 3B, Extended Data Fig. 10), as has been shown for other high efficiency enzymes^32,33^. Enzymes modifying this sabinene backbone to α- and β-thujone are frequently co-inherited as most individuals representing this chemotype contained these monoterpenoids rather than the monoterpene sabinene (Figure 1, Extended Figure S2). Both oxyfunctionalizing enzymes and oxidoreductases were present in the 1.5Mbp specialized metabolism island in the 11-7 genome sequence but not in the syntenic sequence of *A. annua* (Figure 3) suggesting thujone synthesis is localized to a specialized metabolism island. Thujones interact with receptors in the central nervous system and are toxic to both vertebrates and insects^34–36^. Therefore, closely linked enzyme-coding loci for thujone pathways and a highly product-specific MTS are likely selected for. The short evolutionary path to a sabinene synthase and apparent ease of recruitment of modifying enzymes may explain why thujones are present in many other species from diverse lineages (“thujone”, https://roempp.thieme.de/). Membership in different specialized metabolism islands may be the reason for the evolution of a second α-pinene synthase whose locus could not be placed on the 11-7 genome (Figure 2C, 3D, E). In addition to rapid evolution through duplication and neofunctionalization by functional divergence in product spectrum, and finally product specificity, remarkable stability in monoterpenoid production is also observed. The camphene synthases which descended from a common progenitor enzyme retained similar product spectra in both Asteraceae species despite at least 4.5 million years of time since separation^25^ (Figure 3). Tansy individuals dominated by single monoterpenoids like the α-pinene-derived β-chrysanthenyl acetate in line 29-25 support that specificity enhancing molecular architectures such as product-specific MTS enzymes (Figure 2B) with high transcript abundance (Figure 3D, F) are selected for in this species. On the other hand, the local *T. vulgare* population also contained lines with high individual chemodiversity (Figure 1). Enzymes with reduced product specificity can be measured in phylogenetic branches that contain enzymes with high specificity from other lines (Figure 2B). Less specific enzymes can also be generated by *in vitro* mutagenesis of 3-8 amino acids (Figure 3B) underscoring the MTSs known proclivity for broad product spectra with multiple products from a single enzyme^28,29^. In addition to reduced product specificity in enzymes, additional individual chemodiversity enhancing features of terpenoid trait architecture are detected. Present loci of those with presence absence variation (Figure 3C) and the quantitative rather than binary contribution of transcript abundance to product formation (Figure 2D-F) further enhance *individual* chemodiversity (Figure 1). The large changes in monoterpenoids between maternal parent and descendants (Figure 1C-E, Extended Figure S2) suggest large recombination potential which enables population level chemodiversity. The majority of MTS enzymes themselves are indeed unlinked due to the large distances between loci in the 5.1Gbp genome of *T. vulgare* (Figure 2) pointing to a raison d’être for large genomes on one hand and to selection of a genome architecture that enables recombination on the other hand. Expression alleles detected throughout the population with large variation in individual MTS transcript abundances (Figure 2) but surprisingly few pseudogenization events in 11_7 (Extended Figure S11) despite loss of expression in the 11_7 line (Figure 2D) contribute to population level chemodiversity. Biased mutation events outside of gene body regions has been demonstrated in *Arabidopsis thaliana*^97^.

This particular trait architecture with both individual chemodiversity enhancing and reducing features which are readily recombined in the obligate outcrosser resulted in a highly chemodiverse local population of both specific and diverse individual chemotypes. Since the trait architecture enables each individual to be of high or low chemodiversity itself with different dominant monoterpenoids (Figure 1B), each plant contributes to high chemodiversity and therefore patchiness at the local population level. This high chemodiversity of leaf monoterpenoids detected in the local *T. vulgare* population contained in an area within the travel range of pollinators^30^ and large herbivores^31^, was comparable to observations that were made in earlier samplings in similar^2^ and distant^14^ regions (Figure 1A- E, Extended Data Fig. 1 & 2) and therefore is likely a common population level trait. Given that evolution selected for an overall trait architecture that enables population level chemodiversity by selecting for features that enable both low and high individual chemodiversity in closely related individuals, it is feasible to posit the chemodiversity selection hypothesis to explain the vast variation of specialized metabolism observed in nature. In the face of predatory animals capable of association learning, the ability to be different from your neighbors even if you are closely related is apparently important. High chemodiversity at the population level and both high and low chemodiversity at the individual level likely contribute to the success of the invasive weed *T. vulgare* and may provide a blueprint to protect crop monocultures by enhancing a crop’s local population chemodiversity.

## Methods

### *T. vulgare* collection and monoterpenoid characterization

Mature leaves and seeds of *T. vulgare* plants were sampled in January 2019 from a natural population in a ca. 1 km² area in the southern part of Bielefeld, North Rhine-Westphalia, Germany (51°58′–51°59′ N, 8°27′–8°29′ E). Seeds were germinated, grown in the greenhouse and leaf terpenoid profiles of maternal plants and descendants were measured by GC/MS. Twenty-four plants (lines) that were selected for further analysis were clonally propagated by leaf and root cuttings and kept both in the greenhouse and in garden beds near the greenhouse. Sampling locations were visualized in R version 4.5.2^41^ using packages ggmap 4.0.2^43^, sf 1.0- 24^44^ and scatterpie 0.2.6^45^. Map tiles were retrieved from Stadia Maps (Stamen Toner style). Terpenoids were analyzed as described in Ziaja & Müller (2023)^37^. Leaf material was frozen, lyophilized, homogenized, weighed, and extracted in heptane with 1-bromodecane as internal standard in an ultrasonic bath for 5 min. After centrifugation, supernatants were analyzed using gas chromatography coupled with mass spectrometry (GC-MS; GC 2010plus – MS QP2020, Shimadzu, Kyoto, Japan, with VF-5 MS column, 30 m length, 0.2 mm ID, 10 m guard column, Varian, Lake Forest, United States) in electron impact ionization mode at 70 eV and using helium as carrier gas. The starting temperature of 50 °C was kept for 5 min, ramped up to 250 °C at 10 °C min^-1^, increased with 30 °C min^-1^ to a final temperature of 280 °C, and hold for 3 min. Blanks and an alkane standard mix (C7–C40, Sigma Aldrich, Taufkirchen, Germany) were measured under the same conditions. Terpenoids were identified based on their retention indices (RI) and by comparing spectra to synthetic reference compounds, where available, and to entries of the libraries NIST (National Institute of Standards and Technology, Gathersburg, USA, 2014)^38^, Pherobase^39^ and those reported in Adams (2007)^40^. Further GC/MS data analysis and visualization was done in R version 4.5.2^41^ using the packages chemodiv 0.3.1^5^ and ggplot2 4.0.2^42^. Terpenoids in the extracts of the enzyme assays were measured in the same manner.

### Transcriptomes of 24 *T. vulgare* lines

Samples of young leaves of three plant clones of each line were shock-frozen. RNA was isolated using the RNeasy Plant Mini Kit (Qiagen), converted to libraries using the TruSeq Kit (Illumina), and sequenced on the Illumina NextSeq 2000 (not stranded). Two samples per line were sequenced in single-end mode with 70 bp read-length and one sample per line was sequenced in paired-end mode with 2×150 bp read-length. The RNA-Seq data from the 24 lines were quality-trimmed using Trimmomatic v0.39^46^, applying the parameters LEADING:34 TRAILING:34 SLIDINGWINDOW:4:15 ILLUMINACLIP:2:34:15. For single-end read data, a minimum read length of 60 bp was used, while for paired-end data minimum read length was set to 120 bp. Individual *de novo* transcriptome assemblies were generated for each of the 24 *T. vulgare* lines with Trinity v2.15.1^47,48^ based on the trimmed paired-end and single-end RNA-Seq data. Open reading frames were predicted using TransDecoder v5.7.0^49^. The resulting protein sequences were compared to *Arabidopsis thaliana* TAIR10^50^ protein sequences using BLASTP^51^ (BLAST 2.14.0+) and functional annotations transferred. From predicted transcripts, potential TPSs were identified by domain annotation with InterProScan5 v5.68-100.0^52^ using PFAM v35.0^53^ and SUPERFAMILY v1.75^54^ databases. Results were filtered for at least one of the domains PF01397 and PF03936. If several isoforms of the same transcript existed, the longest isoform with both domains was chosen. Chloroplast transit peptides were predicted with TargetP^55^ v2.0 with parameter ‘-org pl’. To place the TPS from *T. vulgare* into an evolutionary context, putative TPS genes and chloroplast transit peptides were identified in *Arabidopsis thaliana* (TAIR10), *Artemisia annua* (<u>GCA_003112345.1</u>), *Marchantia polymorpha* var. BoGa^57,58^, *Cynara cardunculus* var. *scolymus* (<u>GCA_001531365.2</u>)^59^, and *Helianthus annuus* (<u>GCA_002127325.2</u>)^60^ as described above for *T. vulgare* and compared to the longest isoform of every putative TPS gene in *T. vulgare*. All protein sequences were aligned using MUSCLE^61^ v5.1 with the ‘- align’ option. The resulting multiple sequence alignment was used to infer a maximum-likelihood phylogenetic tree with IQ-TREE^62^ v2.3.4 using the parameters ‘-st AA -m TEST - B 1000 -alrt 1000 -T 10 -nm 2000’ and visualized with iTOL^63^ v7.6. Tree branches were assigned to TPS subfamilies according to classified enzymes from *A. thaliana*^64,65^.

### Genome sequencing, assembly, annotation and analysis

Leaf tissue of line 11-7 was sampled from young leaves of healthy plants grown in a garden bed near the greenhouses at Bielefeld University. Leaf material (2.5g) was homogenized in liquid N_2_ with mortar and pestle and incubated in 20 ml extraction buffer (300 mM Tris/HCl, 25 mM EDTA, 2M NaCl, 2% PVP/polyvinylpyrrolidone 40000, 2% CTAB/cetyltrimethylammoniumbromide) for 30 minutes at 65°C. Two times, 1 volume of CIA (24:1 chlorophorm/isoamylalcohol) was added, solution was centrifuged for 15 minutes at 4,500g and supernatant was transferred to a new tube. 1 volume of MilliQ H_2_O was added and solution adjusted to pH 7 (HCl). A genomic tip 100/G column (Quiagen) was equilibrated with 10 ml buffer QBT, extract loaded and incubated until flowed-through by gravity. The column was washed two times with 10 ml buffer QC and DNA eluted with 5 ml buffer QF, preheated to 65°C. 2-propanol (3.5 ml) was added, solution mixed by inverting and incubated over night at 4°C. The solution was centrifuged for 30 minutes (12,000g, 4°C), supernatant removed, pellet washed with 3 ml 80% ethanol, centrifuged for 15 minutes (12,000g, 4°C), supernatant removed and pellet dried at room temperature. DNA was resuspended in 100 µl 10 mM Tris pH8 by incubation at 4°C overnight and subsequent incubations for 15 minutes at 65°C and for 10 minutes on ice. To achieve sufficient genome coverage, genomic DNA (gDNA) from three replicates of *T. vulgare* line 11-7 was sequenced. Sequencing libraries were prepared using the ligation sequencing kit V14 (SQK-LSK114; Oxford Nanopore Technologies (ONT), Oxford, UK) following the manufacturer’s protocol. The initial sample was sequenced on a P2solo device using a single R10.4.1 flow cell. Basecalling was performed with Guppy v6.3.1 using the super-accurate basecalling model (ONT). To increase read length and thus assembly contiguity, two additional gDNA samples from line 11-7 were sequenced. This time, short reads were removed prior to library preparation using the Short Read Eliminator (SRE) XL kit (PacBio), and bases were called with Guppy v6.3.8 (ONT). Adapter sequences were removed using Porechop^66^ v0.2.4. Read correction was performed with Canu^67^ v2.2 using the parameters minReadLength=5,000 and minOverlapLength=3,000. The corrected reads were assembled with Flye^68,69^ v2.9.1, applying the parameters --asm-coverage 30, --genome-size 4.5G, and --nano-corr. The resulting assembly was polished through two rounds of Racon^70^ v1.5.0 followed by two rounds of Medaka v1.8.0 (ONT). Repetitive elements were identified and masked using RepeatModeler^71^ v2.0.4 and RepeatMasker^72^ v4.1.5. Haplotype separation was performed using HaploMerger2^73^ v20180603, resulting in two genome assemblies: one representing the reference haplotype and the other the alternative haplotype. Chromosome-level scaffolding of both haplotypes was carried out with RagTag^74^ v2.1.0, using the *Tanacetum vulgare* genome assembly daTanVulg1.hap1.1 (GenBank accession GCA_964264315.1) as a reference template, and applying the parameters -a 0.1 -i 0.3. Assembly quality and completeness were assessed with BUSCO^75^ v5.8.3 using the eudicotyledons_odb12 database (n=2,805) and QUAST^76^ v5.2.0. Dot plots comparing the two haplotypes and each haplotype with daTanVulg1.hap1.1 were generated using D-Genies^77^ v1.5.0. The haplotype Tv11-7_REF was used for further analyses unless differently specified.

The trimmed RNA-Seq data from the 24 lines were mapped to the reference and to the alternative haplotype with HISAT2^78^ v2.2.1 using the --dta option to optimize for transcript assembly. De novo and reference-guided transcriptome assemblies were computed with Trinity^47,48^ v2.15.2 based on RNA-Seq data from all lines. Paired-end and single-end data were processed separately. A reference-guided assembly was calculated for each haplotype using the parameter --genome_guided_max_intron 10,000. Genes were predicted separately on the reference and alternative haplotype with BRAKER3^79^ v3.0.8 in mode ETP. Paired-end and single-end RNA-Seq alignments served as transcript evidence, and protein homology data from the Viridiplantae dataset^80^ provided additional support. The BUSCO lineage was set to eudicots_odb10. Gene structures were refined with PASA v2.5.3 using the parameters -- ALIGNERS minimap2^81,82^, -N 5, and -I 20000. Additionally, tRNAs were predicted using tRNAscan-SE^83^ v2.0.12.

Protein sequences of the Tv11-7_REF genome were scanned by InterProScan 5.71-102.0^52^ with default parameters and filtered for TPS domains (PF01397 and PF03936). Protein sequences containing at least one out of two domains were annotated by identification of best BlastP^51^ 2.16.0+ matches among TPS candidates identified from transcripts. Candidates with best matches within TPS-b/g subfamily members were used for further analyses. Functions were assigned manually according to the closest experimentally characterized candidate.

Synteny plots were generated with pyGenomeViz 1.6.1^93^ using genomic regions with best matches of characterized sabinene/camphene synthases in *T. vulgare* Tv11-7_REF and *A. annua* GCA_003112345.1^94,95^. Chromosome plots were done with RIdeogram 0.2.2^96^.

### Gene expression analysis

Transcript quantification was performed with Kallisto v0.50.0^56^ by mapping the RNA-Seq reads of each line against each transcriptome and haplotype Tv11-7_REF generated in this study. For single-end reads, the parameters -l 200 -s 20 were applied. Mean TPM values were calculated per line. Expression plots were done with ggplot2 3.5.2^42^ in R version 4.5.2^41^.

To generate product predictions for each MTS, transcript abundance of putative MTS genes were correlated with terpenoid abundance using Spearman’s rank correlation, and correlations with coefficients ≥0.4 were considered significant.

### Protein analyses

Multiple sequence alignments were calculated and visualized in R version 4.5.2^41^ using the msa package^84^ v1.40.0 with ClustalW^85^ 2.1 and TeXshade^86^. Domains and plasticity regions were annotated according to Lei et. al. 2021^87^. Protein structures were predicted using the Chai-1 webserver^88^. Each MTS was modelled with 3 Mg2+ ions and one molecule GPP with standard parameters. The best-scoring model was used for figure generation with ChimeraX^89–91^. Candidate amino acids relevant to product specificity were identified by sequence comparisons between full length sequences of the sabinene and camphene/bornyl diphosphate cluster that evolved from a common ancestor according to the phylogenetic tree generated in this study. Amino acid positions that differed between clusters were considered if they were 100% conserved within the sabinene cluster and at least 85% conserved within the camphene/bornyl diphosphate cluster, analogous to the approach used in Schiller et. al. 2025^92^. Amino acid positions in front of the RRX_8_W motif, that are likely part of the chloroplast targeting peptide were not considered. For each amino acid position, the literature was checked whether the amino acid at that position is known to be involved in substrate binding or to be part of the plasticity regions known to affect the product outcome in other species. The amino acids were located in the 3D protein structure predicted by Chai-1^88^ and proximity to the substrate in the binding pocket was used for selection of positions for mutation.

### Enzyme assays

Transcript sequences encoding the mature proteins starting from the first conserved ‘RR’ residues were cloned with an leading methionine into the pET16b expression vector using the BamHI and NdeI restriction sites. If none or multiple ‘RR’ were found, chloroplast transit peptides of selected MTS candidate transcripts were predicted using TargetP2.0^55^ (htMTS://services.healthtech.dtu.dk/services/TargetP-2.0/, ‘plant’ setting) to determine the sequences to omit. Inserted sequences were validated by Sanger sequencing. Gibson primers and insert sequences are listed in Supplemental file 2. For protein expression, cultures (TBY media with 100µg/mL ampicillin) were inoculated from overnight cultures with an OD600 of 0.1 and grown at 37°C to an OD600 of 0.6-0.8. 0.5 mM IPTG was added and cultivation flasks were incubated at 22°C overnight. Cells were harvested by centrifugation at 4,000 rpm for 20 minutes at 4°C. Pellets were stored at -20°C until the measurements were performed. For the enzyme assay, pellets were thawed on ice and resuspended in 3 or 6 mL/g assay buffer (50 mM PIPES, pH 7.6, 10 mM MgCl2, 100 mM NaCl, 2 mM DTT), lysed using sonication and centrifuged at 4,000 rpm for 20 minutes at 4°C. To the supernatant, 300 µM GPP (in 25 mM NH4HCO3) was added, the mixture was overlayed with one volume heptane and incubated for 2h at room temperature. The reaction was stopped by vortexing. After phases were separated again, the heptane phase was transferred with Pasteur pipettes to glass vials for GC/MS measurements which were performed analogous to leaf terpenoid measurements. Mean peak areas of control (empty vector) samples were subtracted from other samples to account for trace compounds from *E. coli* lysates.

### LLM use

Perplexity AI (Grok 4.1, accessed September 2025–May 2026) was used to retrieve literature, verify citations, assist text editing and generate code snippets for R and Python. Code was manually adapted, tested, and validated by authors. All factual claims and code outputs were manually verified and executed on local clusters. All retrieved information was manually validated against primary sources. Authors assume full responsibility.

## Data availability

The annotated genome haplotypes are available under PRJNA1481999 and PRJNA1481998, respectively. The transcriptome data has been deposited under PRJEB115532.

## Acknowledgements

The authors thank Sebastian Tschikin for RNA sampling of the 24 *Tanacetum vulgare* lines, Christine Schlüter and the Bielefeld University greenhouse team for plant care, the NGS team of the Bielefeld University Omics CF NGS Unit and CeBiTec as well as the technical staff of the CeBiTec Technology Platform Genomics for technical assistance, the Bioinformatics Resource Facility for Compute and Storage Infrastructure, and Dr. Kai Schülke for advice on the enzyme assays

Financial support by Deutsche Forschungsgemeinschaft to CM (FOR3000, project number 415496540 and MU1829/28-2) and to AB (BR4627-1, BR4627-2) is gratefully acknowledged.

## Contributions

MH analyzed the transcriptomes and genome, functionally tested the enzymes, performed the evolutionary analyses and co-wrote the manuscript; BL assembled and annotated the genome and transcriptomes and performed phylogenetic analyses; EE collected the plants and EE, DZ, and RJ performed GC-MS measurements; PV performed transcriptome sequencing; TB supervised genome sequencing; SH supported the reconstruction of terpenoid synthesis pathways; ME supported enzyme assays; CM obtained funding and advised on the analyses; AB obtained funding, advised on the analyses, developed the concepts, and co-wrote the manuscript; all authors commented on and edited the manuscript

## References

1. Weng, J.-K., Lynch, J. H., Matos, J. O. & Dudareva, N. Adaptive mechanisms of plant specialized metabolism connecting chemistry to function. Nat. Chem. Biol. 17, 1037–1045 (2021).

2. Kleine, S. & Müller, C. Intraspecific plant chemical diversity and its relation to herbivory. Oecologia 166, 175–186 (2011).

3. Ziaja, D., Sasidharan, R., Jakobs, R., Eilers, E. J. & Müller, C. Chemotype, maternal genotype, or field neighbors: what influences performance and resource allocation in a perennial plant species the most? Oecologia 207, 134 (2025).

4. Sasidharan, R., Grond, S. G., Champion, S., Eilers, E. J. & Müller, C. Intraspecific plant chemodiversity at the individual and plot levels influences flower visitor groups with consequences for germination success. Funct. Ecol. 1365-2435.14673 (2024) doi:10.1111/1365-2435.14673.

5. Petrén, H., Köllner, T. G. & Junker, R. R. Quantifying chemodiversity considering biochemical and structural properties of compounds with the R package CHEMODIV. New Phytol. 237, 2478–2492 (2023).

6. Petrén, H. et al. Understanding the chemodiversity of plants: Quantification, variation and ecological function. Ecol. Monogr. 94, e1635 (2024).

7. Wetzel, W. C. & Whitehead, S. R. The many dimensions of phytochemical diversity: linking theory to practice. Ecol. Lett. 23, 16–32 (2020).

8. Thon, F. M., Müller, C. & Wittmann, M. J. The evolution of chemodiversity in plants— From verbal to quantitative models. Ecol. Lett. 27, e14365 (2024).

9. Wittmann, M. J. & Bräutigam, A. How does plant chemodiversity evolve? Testing five hypotheses in one population genetic model. New Phytol. nph.20096 (2024) doi:10.1111/nph.20096.

10. Katz, E. et al. Genetic variation, environment and demography intersect to shape Arabidopsis defense metabolite variation across Europe. eLife 10, e67784 (2021).

11. Lawrence, E. J., Griffin, C. H. & Henderson, I. R. Modification of meiotic recombination by natural variation in plants. J. Exp. Bot. 68, 5471–5483 (2017).

12. Zhou, X. & Liu, Z. Unlocking plant metabolic diversity: A (pan)-genomic view. Plant Commun. 3, 100300 (2022).

13. Richards, C. L., Bossdorf, O., Muth, N. Z., Gurevitch, J. & Pigliucci, M. Jack of all trades, master of some? On the role of phenotypic plasticity in plant invasions. Ecol. Lett. 9, 981–993 (2006).

14. Wolf, V. C., Berger, U., Gassmann, A. & Müller, C. High chemical diversity of a plant species is accompanied by increased chemical defence in invasive populations. Biol. Invasions 13, 2091–2102 (2011).

15. Ojeda Prieto, L., Medina van Berkum, P., Unsicker, S. B., Heinen, R. & Weisser, W. W. Intraspecific chemical variation of *Tanacetum vulgare* affects plant growth and reproductive traits in field plant communities. Plant Biol. 27, 785–801 (2025).

16. Ziaja, D. & Müller, C. Intraspecific and intra individual chemodiversity and phenotypic integration of terpenes across plant parts and development stages in an aromatic plant. Plant Biol. 27, 637–650 (2025).

17. Holopainen, M., Hiltunen, R. & Von Schantz, M. A Study on Tansy Chemotypes. Planta Med. 53, 284–287 (1987).

18. Müller, C., Dussarrat, T. & van Dam, N. M. What is a plant chemotype anyway? Trends Plant Sci. (2026).

19. Srividya, N., Kim, H., Simone, R. & Lange, B. M. Chemical diversity in angiosperms − monoterpene synthases control complex reactions that provide the precursors for ecologically and commercially important monoterpenoids. Plant J. 119, 28–55 (2024).

20. Brown, G. D. The Biosynthesis of Artemisinin (Qinghaosu) and the Phytochemistry of Artemisia annua L. (Qinghao). Molecules 15, 7603–7698 (2010).

21. Banthorpe, D. V., Doonan, S. & Gutowski, J. A. Biosynthesis of irregular monoterpenes in extracts from higher plants. Phytochemistry 16, 85–92 (1977).

22. Jiang, S.-Y., Jin, J., Sarojam, R. & Ramachandran, S. A Comprehensive Survey on the Terpene Synthase Gene Family Provides New Insight into Its Evolutionary Patterns. Genome Biol. Evol. 11, 2078–2098 (2019).

23. Darwin Tree of Life Project. Tanacetum vulgare genome assembly GCA_964264315.1, INSDC ID PRJEB74721. https://portal.darwintreeoflife.org/data/Tanacetum%20vulgare [20.03.2025].

24. Ruan, J.-X. et al. Isolation and Characterization of Three New Monoterpene Synthases from Artemisia annua. Front. Plant Sci. 7, (2016).

25. Kumar, S., Stecher, G., Suleski, M. & Hedges, S. B. TimeTree: A Resource for Timelines, Timetrees, and Divergence Times. Mol. Biol. Evol. 34, 1812–1819 (2017).

26. Weng, J.-K., Philippe, R. N. & Noel, J. P. The Rise of Chemodiversity in Plants. Science 336, 1667–1670 (2012).

27. Ono, E. & Murata, J. Exploring the Evolvability of Plant Specialized Metabolism: Uniqueness Out Of Uniformity and Uniqueness Behind Uniformity. Plant Cell Physiol. 64, 1482–1493 (2023).

28. Whitehead, J. N., Leferink, N. G. H., Johannissen, L. O., Hay, S. & Scrutton, N. S. Decoding Catalysis by Terpene Synthases. ACS Catal. 13, 12774–12802 (2023).

29. Yoshikuni, Y., Ferrin, T. E. & Keasling, J. D. Designed divergent evolution of enzyme function. Nature 440, 1078–1082 (2006).

30. Greenleaf, S. S., Williams, N. M., Winfree, R. & Kremen, C. Bee foraging ranges and their relationship to body size. Oecologia 153, 589–596 (2007).

31. De Knegt, H. J., Hengeveld, G. M., Van Langevelde, F., De Boer, W. F. & Kirkman, K. P. Patch density determines movement patterns and foraging efficiency of large herbivores. Behav. Ecol. 18, 1065–1072 (2007).

32. Hunt, S. E. et al. Distal mutations in a designed retro-aldolase alter loop dynamics to shift and accelerate the rate-limiting step. Preprint at 10.1101/2025.01.26.634918 (2025).

33. Romero-Rivera, A., Garcia-Borràs, M. & Osuna, S. Role of Conformational Dynamics in the Evolution of Retro-Aldolase Activity. ACS Catal. 7, 8524–8532 (2017).

34. Höld, K. M., Sirisoma, N. S., Ikeda, T., Narahashi, T. & Casida, J. E. α-Thujone (the active component of absinthe): γ-Aminobutyric acid type A receptor modulation and metabolic detoxification. Proc. Natl. Acad. Sci. 97, 3826–3831 (2000).

35. Deiml, T. et al. α-Thujone reduces 5-HT3 receptor activity by an effect on the agonist-induced desensitization. Neuropharmacology 46, 192–201 (2004).

36. Meschler, J. Thujone Exhibits Low Affinity for Cannabinoid Receptors But Fails to Evoke Cannabimimetic Responses. Pharmacol. Biochem. Behav. 62, 473–480 (1999).

37. Ziaja, D. & Müller, C. Intraspecific chemodiversity provides plant individual- and neighbourhood-mediated associational resistance towards aphids. Front. Plant Sci. 14, 1145918 (2023).

38. National Institute of Standards and Technology. NIST/EPA/NIH Mass Spectral Library. (2014).

39. El-Sayed, A. M. The Pherobase: Database of Pheromones and Semiochemicals. (2012).

40. Adams, R. P. Identification of Essential Oil Components by Gas Chromatography/Quadrupole Mass Spectroscopy. (Allured Publishing Corporation, 2007).

41. R Core Team. R: A Language and Environment for Statistical Computing. R Foundation for Statistical Computing (2026).

42. Wickham, H. Ggplot2: Elegant Graphics for Data Analysis. (Springer International Publishing : Imprint: Springer, Cham, 2016). doi:10.1007/978-3-319-24277-4.

43. Kahle, D. & Wickham, H. Ggmap: Spatial Visualization with Ggplot2. (2025).

44. Pebesma, E. Sf: Simple Features for R. (2026).

45. Yu, G. Scatterpie: Scatter Pie Plot. (2026). doi:10.32614/CRAN.package.scatterpie.

46. Bolger, A. M., Lohse, M. & Usadel, B. Trimmomatic: a flexible trimmer for Illumina sequence data. Bioinformatics vol. 30 2114–2120 (2014).

47. Haas, B. J. et al. De novo transcript sequence reconstruction from RNA-seq using the Trinity platform for reference generation and analysis. Nat. Protoc. 8, 1494–1512 (2013).

48. Grabherr, M. G., et al. Full-length transcriptome assembly from RNA-Seq data without a reference genome. (2011).

49. Haas, B. J. TransDecoder. (2019).

50. The Arabidopsis Information Resource. Arabidopsis thaliana genome assembly TAIR10. (2011).

51. Camacho, C. et al. BLAST+: architecture and applications. BMC Bioinformatics 10, 421 (2009).

52. Jones, P. et al. InterProScan 5: genome-scale protein function classification. Bioinformatics 30, 1236–1240 (2014).

53. Paysan-Lafosse, T. et al. The Pfam protein families database: embracing AI/ML. Nucleic Acids Res. 53, D523–D534 (2025).

54. Oates, M. E. et al. The SUPERFAMILY 1.75 database in 2014: a doubling of data. Nucleic Acids Res. 43, D227–D233 (2015).

55. Almagro Armenteros, J. J., et al. Detecting sequence signals in targeting peptides using deep learning. Life Sci. Alliance 2, e201900429 (2019).

56. Bray, N. L., Pimentel, H., Melsted, P. & Pachter, L. Near-optimal probabilistic RNA-seq quantification. Nat. Biotechnol. 34, 525–527 (2016).

57. Frommer, B. & Bräutigam, A. Genome sequence assembly of the liverwort Marchantia polymorpha subsp. ruderalis ecotype BoGa and its gene annotation. 70269095 bytes, 11134555 bytes Bielefeld University 10.4119/UNIBI/2982437 (2023).

58. Beaulieu, C. et al. The Marchantia polymorpha pangenome reveals ancient mechanisms of plant adaptation to the environment. Nat. Genet. 57, 729–740 (2025).

59. Scaglione, D. et al. The genome sequence of the outbreeding globe artichoke constructed de novo incorporating a phase-aware low-pass sequencing strategy of F1 progeny. Sci. Rep. 6, 19427 (2016).

60. Badouin, H. et al. The sunflower genome provides insights into oil metabolism, flowering and Asterid evolution. Nature 546, 148–152 (2017).

61. Edgar, R. C. Muscle5: High-accuracy alignment ensembles enable unbiased assessments of sequence homology and phylogeny. Nat. Commun. 13, 6968 (2022).

62. Minh, B. Q. et al. IQ-TREE 2: New Models and Efficient Methods for Phylogenetic Inference in the Genomic Era. Mol. Biol. Evol. 37, 1530–1534 (2020).

63. Letunic, I. & Bork, P. Interactive Tree of Life (iTOL) v6: recent updates to the phylogenetic tree display and annotation tool. Nucleic Acids Res. 52, W78–W82 (2024).

64. Parker, M. T., Zhong, Y., Dai, X., Wang, S. & Zhao, P. Comparative genomic and transcriptomic analysis of terpene synthases in *Arabidopsis* and *Medicago*. IET Syst. Biol. 8, 146–153 (2014).

65. Aubourg, S., Lecharny, A. & Bohlmann, J. Genomic analysis of the terpenoid synthase (AtTPS) gene family of Arabidopsis thaliana. Mol. Genet. Genomics 267, 730– 745 (2002).

66. Wick, R. R., Judd, L. M., Gorrie, C. L. & Holt, K. E. Completing bacterial genome assemblies with multiplex MinION sequencing. *Microb*. Genomics 3, (2017).

67. Koren, S. et al. Canu: scalable and accurate long-read assembly via adaptive *k* -mer weighting and repeat separation. Genome Res. 27, 722–736 (2017).

68. Lin, Y. et al. Assembly of long error-prone reads using de Bruijn graphs. Proc. Natl. Acad. Sci. 113, (2016).

69. Kolmogorov, M., Yuan, J., Lin, Y. & Pevzner, P. A. Assembly of long, error-prone reads using repeat graphs. Nat. Biotechnol. 37, 540–546 (2019).

70. Vaser, R., Sović, I., Nagarajan, N. & Šikić, M. Fast and accurate de novo genome assembly from long uncorrected reads. Genome Res. 27, 737–746 (2017).

71. Flynn, J. M. et al. RepeatModeler2 for automated genomic discovery of transposable element families. Proc. Natl. Acad. Sci. 117, 9451–9457 (2020).

72. Smit, A., Hubley, R. & Green, P. RepeatMasker Open-4.0. (2013).

73. Huang, S., Kang, M. & Xu, A. HaploMerger2: rebuilding both haploid sub-assemblies from high-heterozygosity diploid genome assembly. Bioinformatics 33, 2577–2579 (2017).

74. Alonge, M. et al. Automated assembly scaffolding using RagTag elevates a new tomato system for high-throughput genome editing. Genome Biol. 23, 258 (2022).

75. Manni, M., Berkeley, M. R., Seppey, M., Simão, F. A. & Zdobnov, E. M. BUSCO Update: Novel and Streamlined Workflows along with Broader and Deeper Phylogenetic Coverage for Scoring of Eukaryotic, Prokaryotic, and Viral Genomes. Mol. Biol. Evol. 38, 4647–4654 (2021).

76. Mikheenko, A., Prjibelski, A., Saveliev, V., Antipov, D. & Gurevich, A. Versatile genome assembly evaluation with QUAST-LG. Bioinformatics 34, i142–i150 (2018).

77. Cabanettes, F. & Klopp, C. D-GENIES: dot plot large genomes in an interactive, efficient and simple way. PeerJ 6, e4958 (2018).

78. Kim, D., Paggi, J. M., Park, C., Bennett, C. & Salzberg, S. L. Graph-based genome alignment and genotyping with HISAT2 and HISAT-genotype. Nat. Biotechnol. 37, 907– 915 (2019).

79. Gabriel, L. et al. BRAKER3: Fully automated genome annotation using RNA-seq and protein evidence with GeneMark-ETP, AUGUSTUS, and TSEBRA. Genome Res. 34, 769– 777 (2024).

80. Tegenfeldt, F. et al. OrthoDB and BUSCO update: annotation of orthologs with wider sampling of genomes. Nucleic Acids Res. 53, D516–D522 (2025).

81. Haas, B. J. Improving the Arabidopsis genome annotation using maximal transcript alignment assemblies. Nucleic Acids Res. 31, 5654–5666 (2003).

82. Campbell, M. A., Haas, B. J., Hamilton, J. P., Mount, S. M. & Buell, C. R. Comprehensive analysis of alternative splicing in rice and comparative analyses with Arabidopsis. BMC Genomics 7, 327 (2006).

83. Chan, P. P., Lin, B. Y., Mak, A. J. & Lowe, T. M. tRNAscan-SE 2.0: improved detection and functional classification of transfer RNA genes. Nucleic Acids Res. 49, 9077– 9096 (2021).

84. Bodenhofer, U., Bonatesta, E., Horejš-Kainrath, C. & Hochreiter, S. msa: an R package for multiple sequence alignment. Bioinformatics 31, 3997–3999 (2015).

85. Thompson, J. D., Higgins, D. G. & Gibson, T. J. Clustal W: improving the sensitivity of progressive multiple sequence alignment through sequence weighting, position-specific gap penalties and weight matrix choice.

86. Beitz, E. TEXshade: shading and labeling of multiple sequence alignments using LATEX 2ε.

87. Lei, D., Qiu, Z., Qiao, J. & Zhao, G.-R. Plasticity engineering of plant monoterpene synthases and application for microbial production of monoterpenoids. Biotechnol. Biofuels 14, 147 (2021).

88. Chai Discovery et al. Chai-1: Decoding the molecular interactions of life. Preprint at 10.1101/2024.10.10.615955 (2024).

89. Goddard, T. D. et al. UCSF ChimeraX: Meeting modern challenges in visualization and analysis. Protein Sci. 27, 14–25 (2018).

90. Meng, E. C. et al. UCSF CHIMERAX : Tools for structure building and analysis. Protein Sci. 32, e4792 (2023).

91. Pettersen, E. F. et al. UCSF CHIMERAX : Structure visualization for researchers, educators, and developers. Protein Sci. 30, 70–82 (2021).

92. Schiller, K. Regulation of Crassulacean acid metabolism at the protein level in Kalanchoë laxiflora.

93. Shimoyama, Y. pyGenomeViz: A genome visualization python package for comparative genomics. (2024).

94. NCBI Genome Dataset. Artemisia annua Huhao1 genome assembly GCA_003112345.1 (ASM311234v1). (2018).

95. Shen, Q. et al. The Genome of Artemisia annua Provides Insight into the Evolution of Asteraceae Family and Artemisinin Biosynthesis. Mol. Plant 11, 776–788 (2018).

96. Hao, Z., et al. *RIdeogram* : drawing SVG graphics to visualize and map genome-wide data on the idiograms. PeerJ Comput. Sci. 6, e251 (2020).

97. Monroe, J.G., Srikant, T., Carbonell-Bejerano, P. et al. Mutation bias reflects natural selection in Arabidopsis thaliana. Nature 602, 101–105 (2022).

