## Supplemental Figures for "Molecular sources of monoterpenoid chemodiversity in the Asteraceae *Tanacetum vulgare* suggest a new model for the evolution of specialized metabolism"

### Slide 1
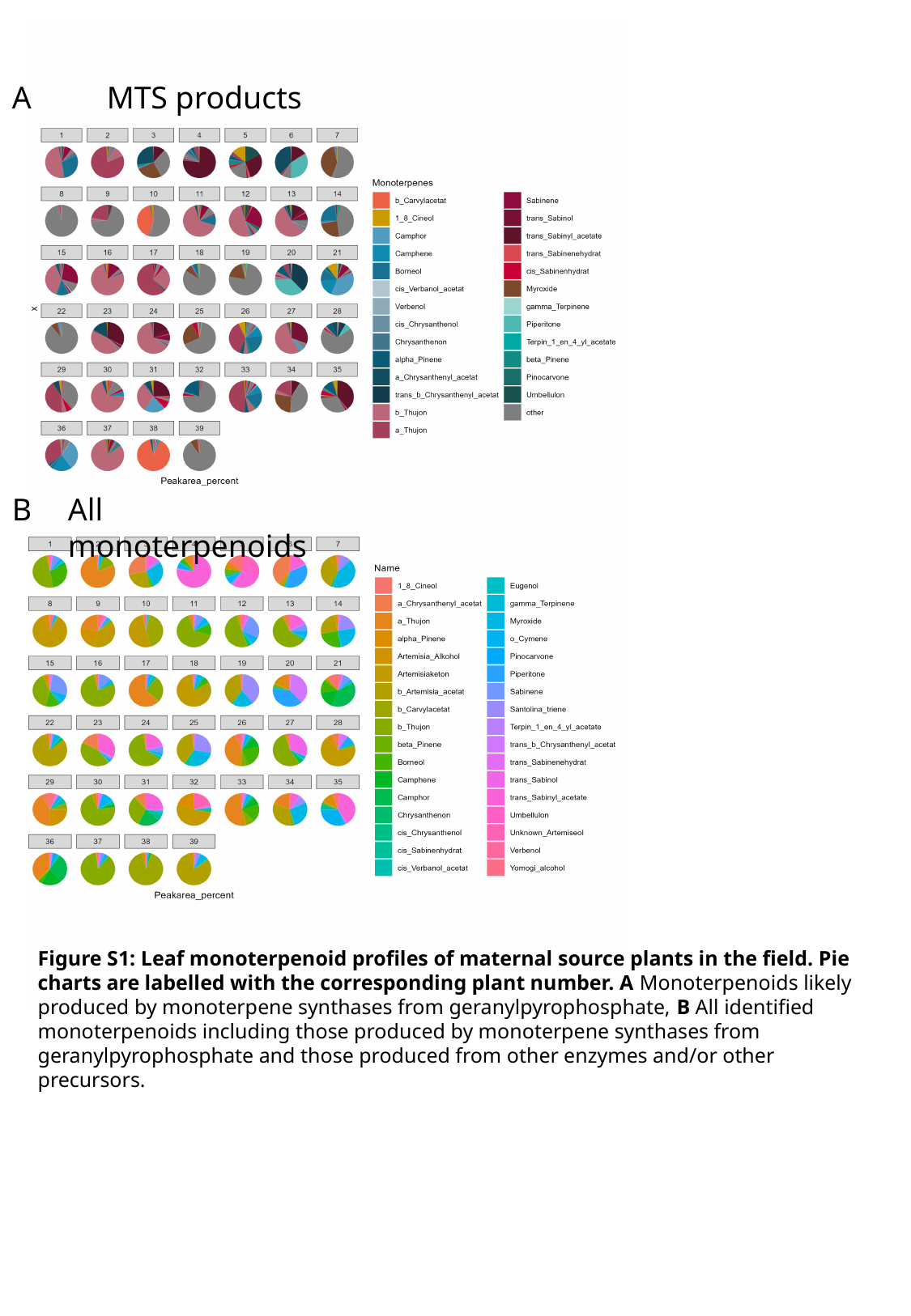

MTS products
A
B
All monoterpenoids
Figure S1: Leaf monoterpenoid profiles of maternal source plants in the field. Pie charts are labelled with the corresponding plant number. A Monoterpenoids likely produced by monoterpene synthases from geranylpyrophosphate, B All identified monoterpenoids including those produced by monoterpene synthases from geranylpyrophosphate and those produced from other enzymes and/or other precursors.

### Slide 2
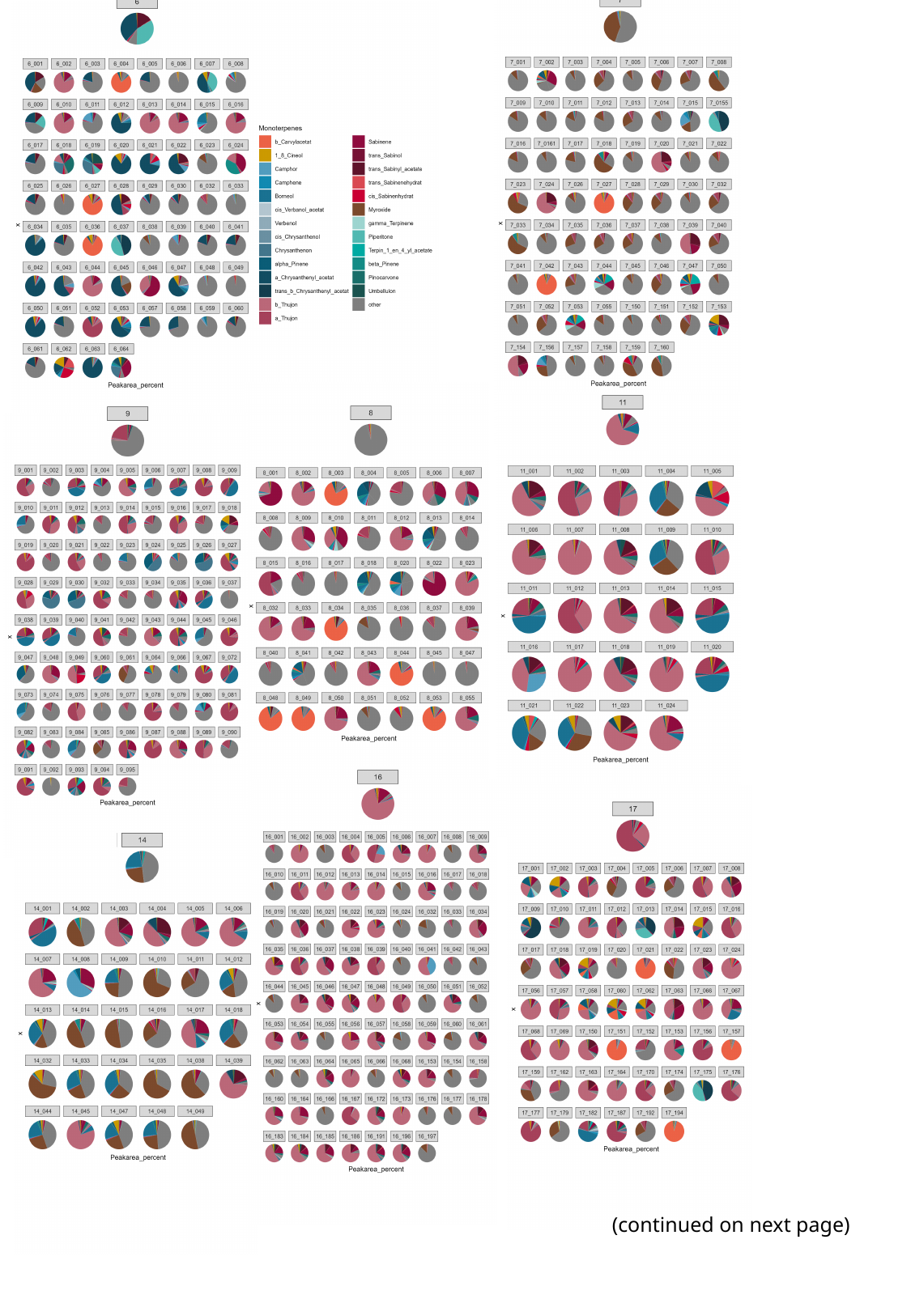

(continued on next page)

### Slide 3
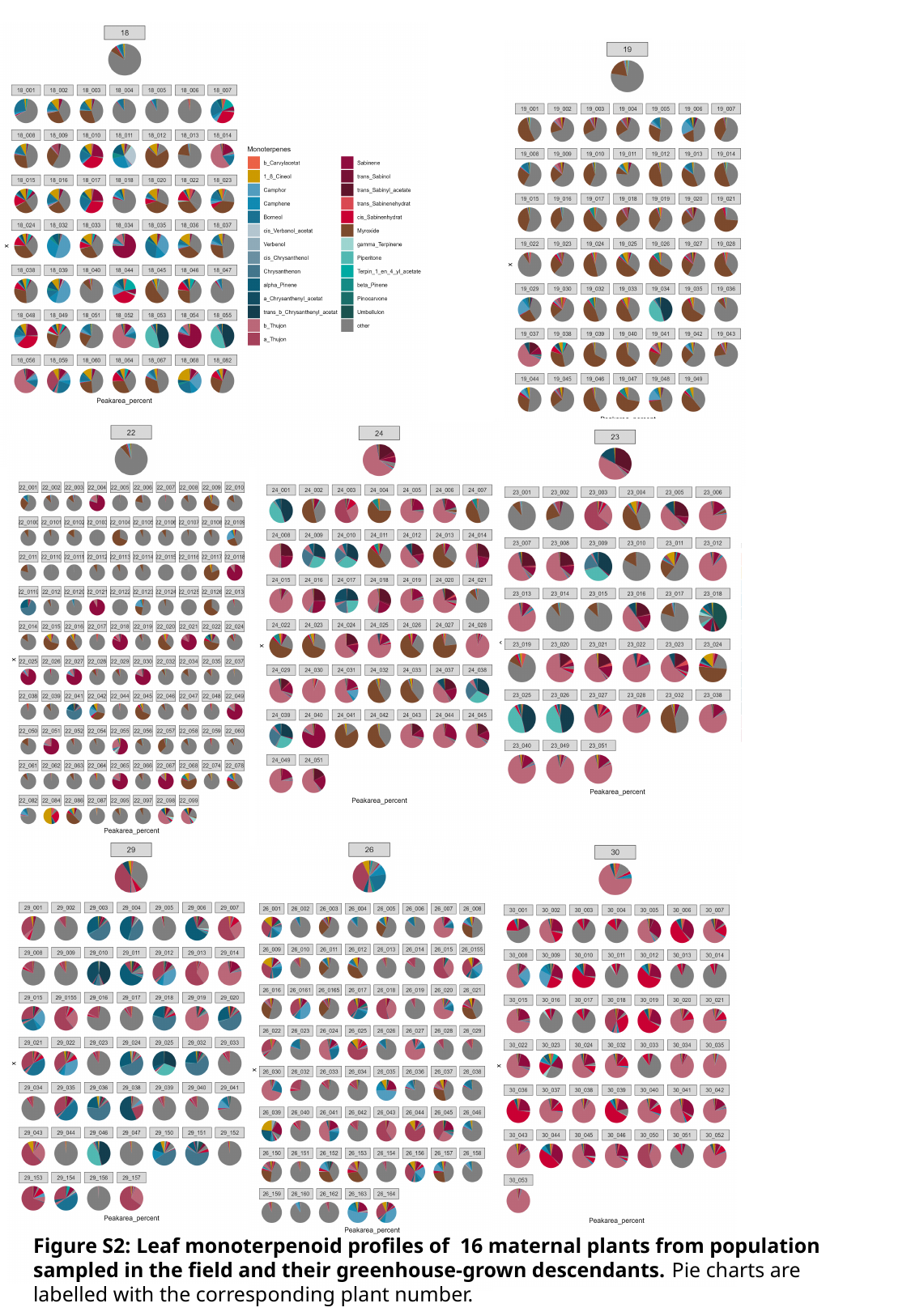

Figure S2: Leaf monoterpenoid profiles of 16 maternal plants from population sampled in the field and their greenhouse-grown descendants. Pie charts are labelled with the corresponding plant number.

### Slide 4
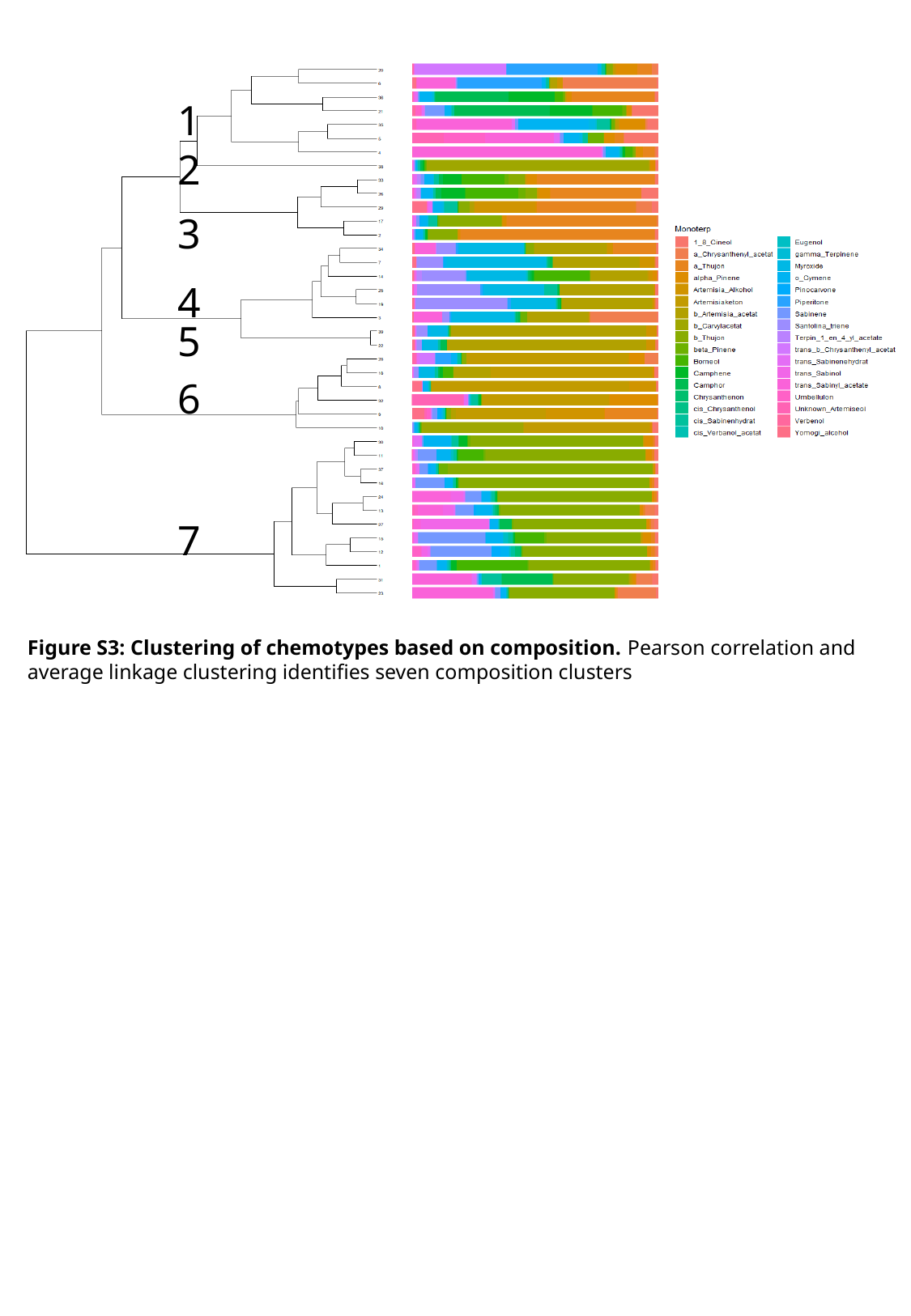

1
2
3
4
5
6
7
Figure S3: Clustering of chemotypes based on composition. Pearson correlation and average linkage clustering identifies seven composition clusters

### Slide 5
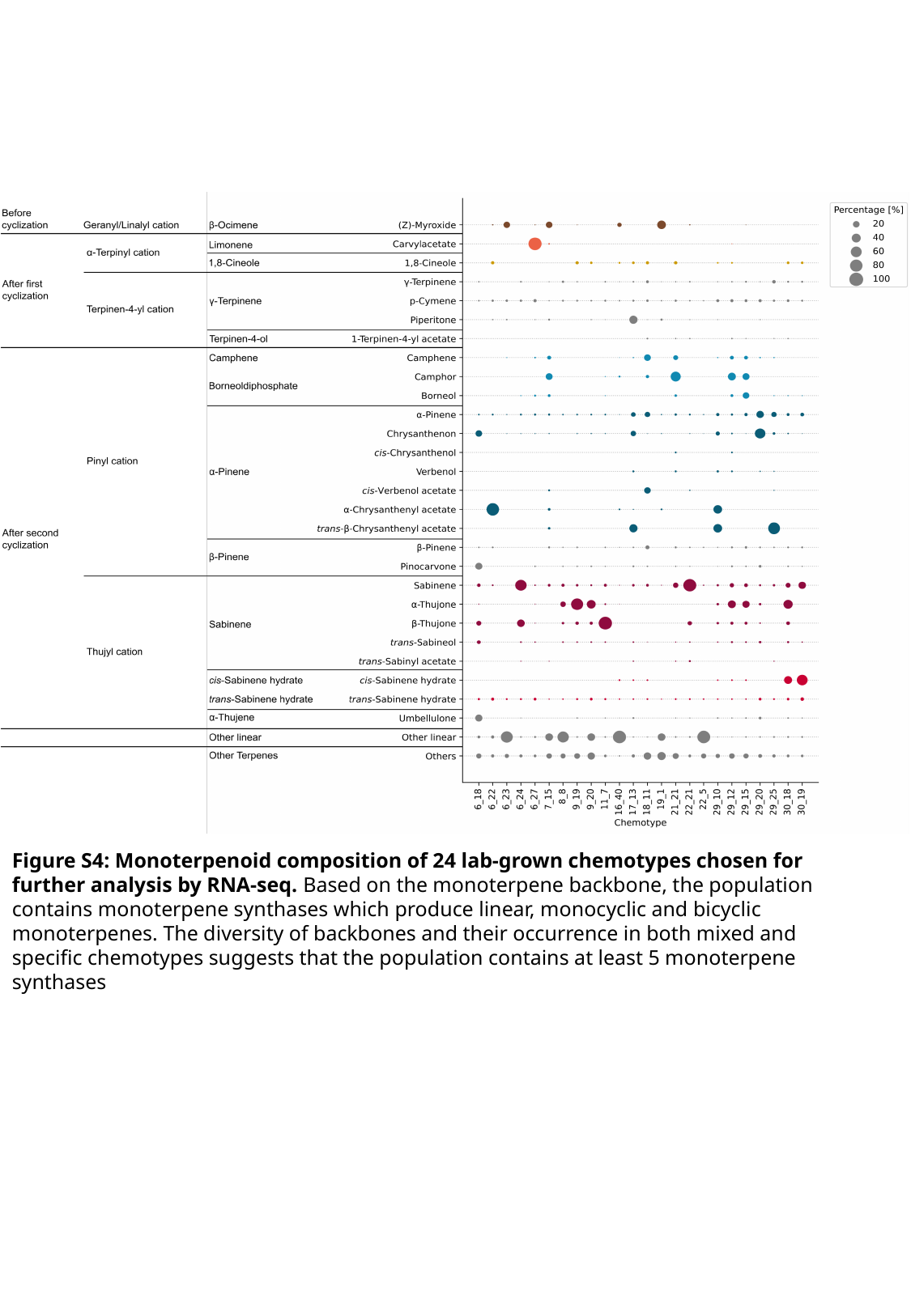

Figure S4: Monoterpenoid composition of 24 lab-grown chemotypes chosen for further analysis by RNA-seq. Based on the monoterpene backbone, the population contains monoterpene synthases which produce linear, monocyclic and bicyclic monoterpenes. The diversity of backbones and their occurrence in both mixed and specific chemotypes suggests that the population contains at least 5 monoterpene synthases

### Slide 6
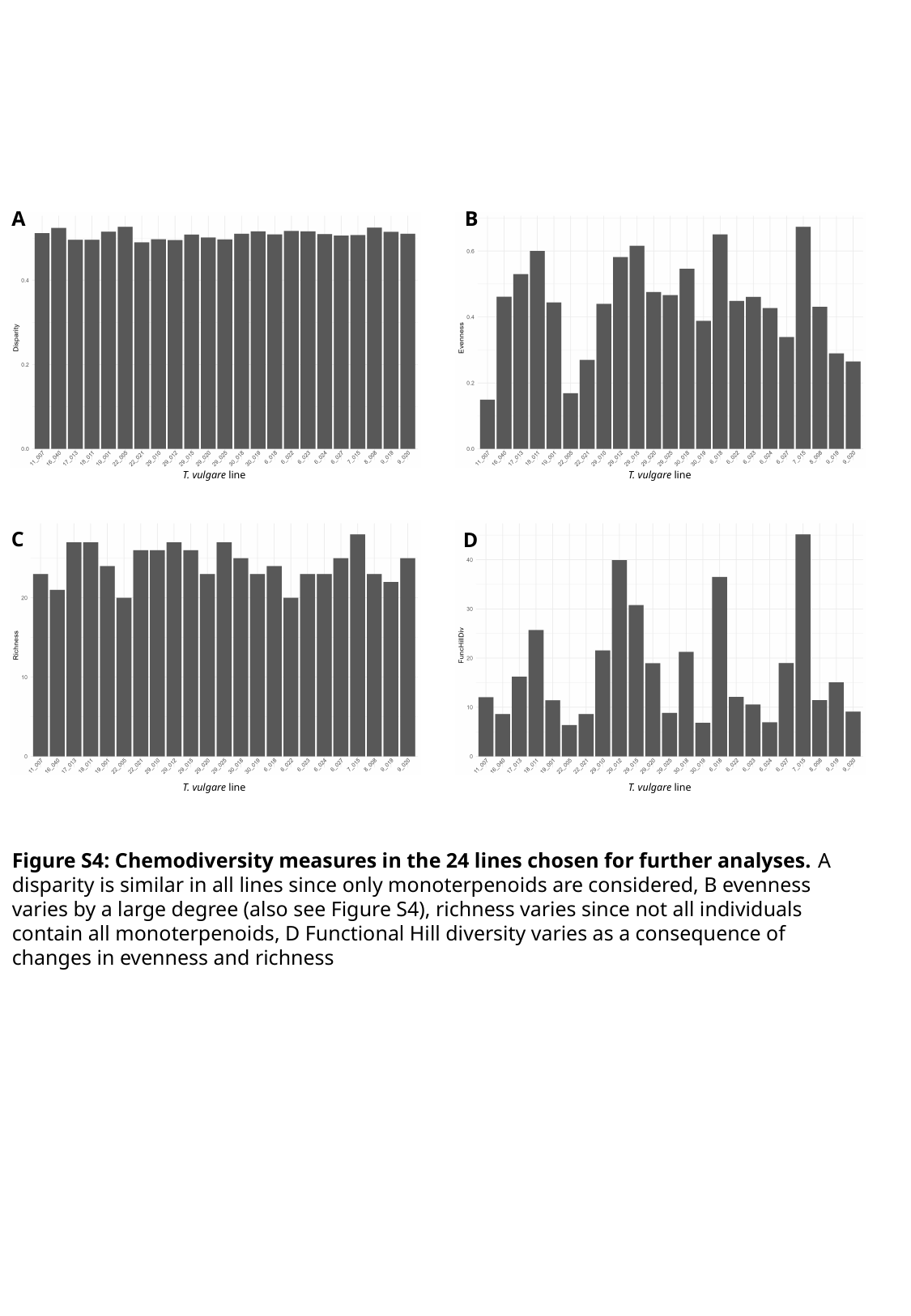

A
B
T. vulgare line
T. vulgare line
C
D
T. vulgare line
T. vulgare line
Figure S4: Chemodiversity measures in the 24 lines chosen for further analyses. A disparity is similar in all lines since only monoterpenoids are considered, B evenness varies by a large degree (also see Figure S4), richness varies since not all individuals contain all monoterpenoids, D Functional Hill diversity varies as a consequence of changes in evenness and richness

### Slide 7
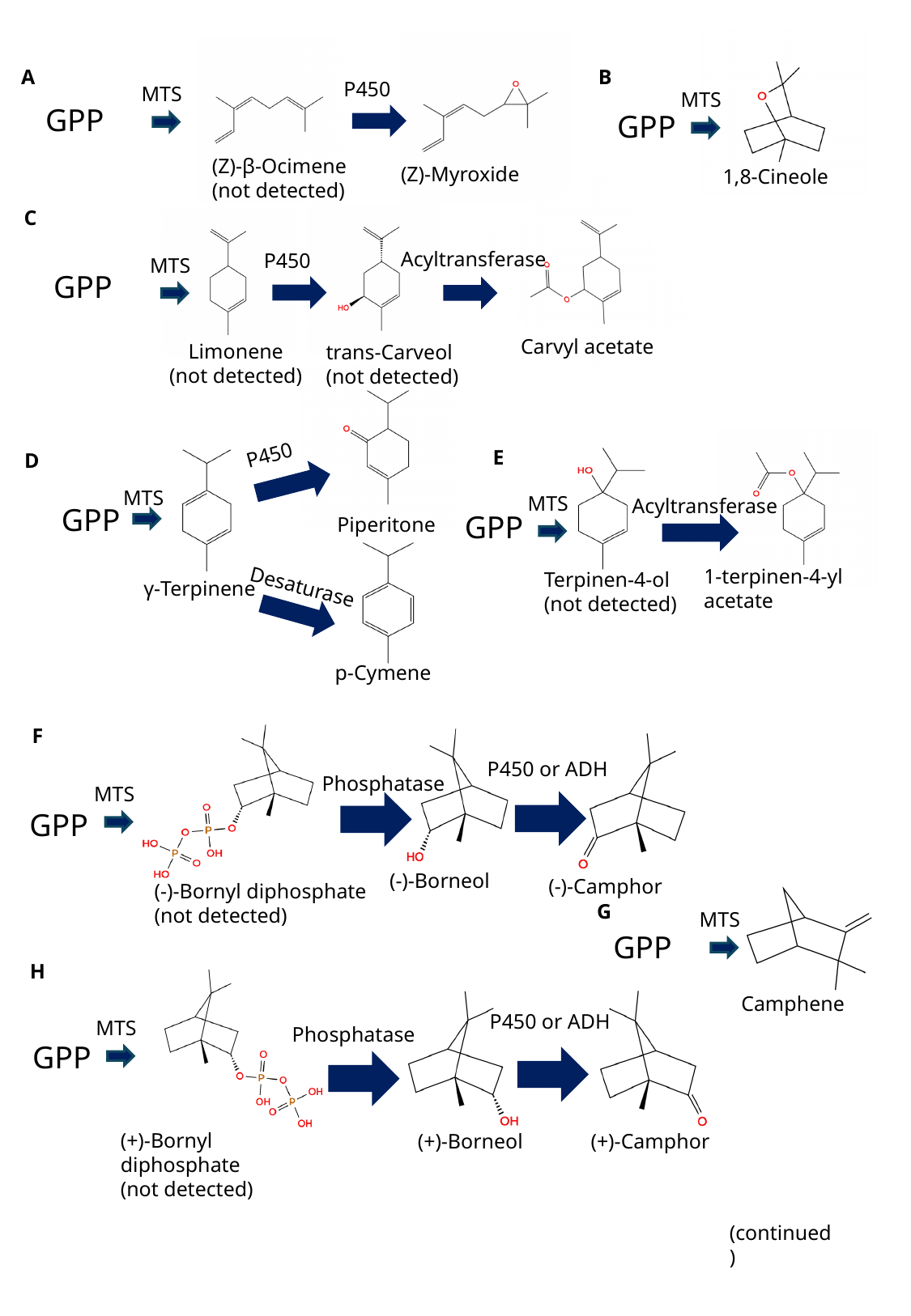

MTS
1,8-Cineole
(Z)-Myroxide
(Z)-β-Ocimene
(not detected)
A
B
P450
MTS
GPP
GPP
Carvyl acetate
C
Limonene
(not detected)
Acyltransferase
P450
MTS
GPP
trans-Carveol
(not detected)
Piperitone
γ-Terpinene
P450
MTS
GPP
p-Cymene
Desaturase
MTS
Acyltransferase
GPP
Terpinen-4-ol
(not detected)
1-terpinen-4-yl acetate
E
D
(-)-Bornyl diphosphate
(not detected)
(-)-Borneol
(-)-Camphor
F
P450 or ADH
Phosphatase
MTS
GPP
MTS
GPP
Camphene
G
(+)-Bornyl diphosphate
(not detected)
H
(+)-Camphor
(+)-Borneol
P450 or ADH
MTS
Phosphatase
GPP
(continued)

### Slide 8
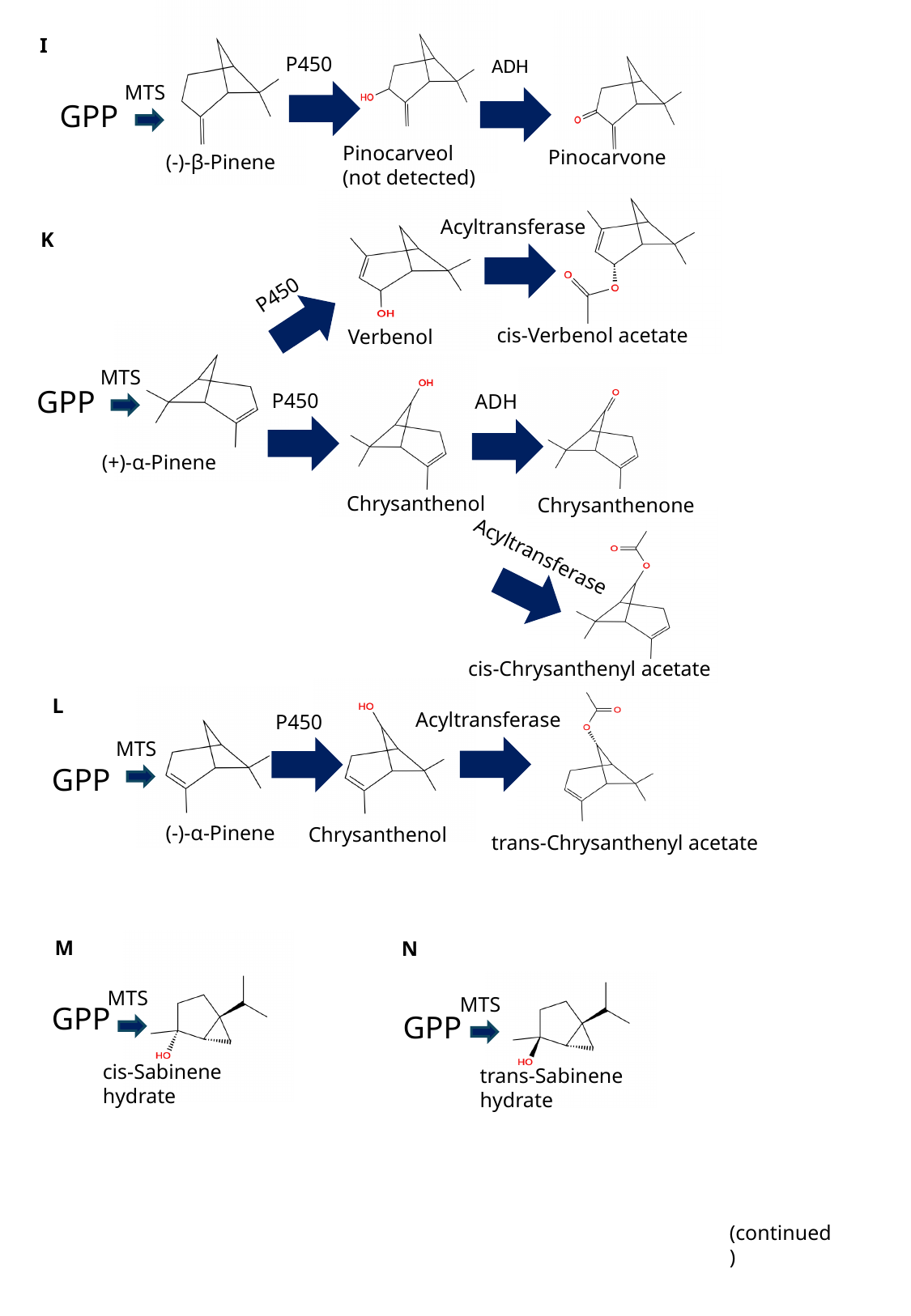

Pinocarveol
(not detected)
(-)-β-Pinene
Pinocarvone
I
P450
ADH
MTS
GPP
cis-Verbenol acetate
Verbenol
Acyltransferase
K
P450
(+)-α-Pinene
Chrysanthenol
MTS
Chrysanthenone
GPP
P450
ADH
cis-Chrysanthenyl acetate
Acyltransferase
trans-Chrysanthenyl acetate
(-)-α-Pinene
L
Acyltransferase
P450
MTS
GPP
Chrysanthenol
M
N
MTS
cis-Sabinene hydrate
GPP
MTS
trans-Sabinene hydrate
GPP
(continued)

### Slide 9
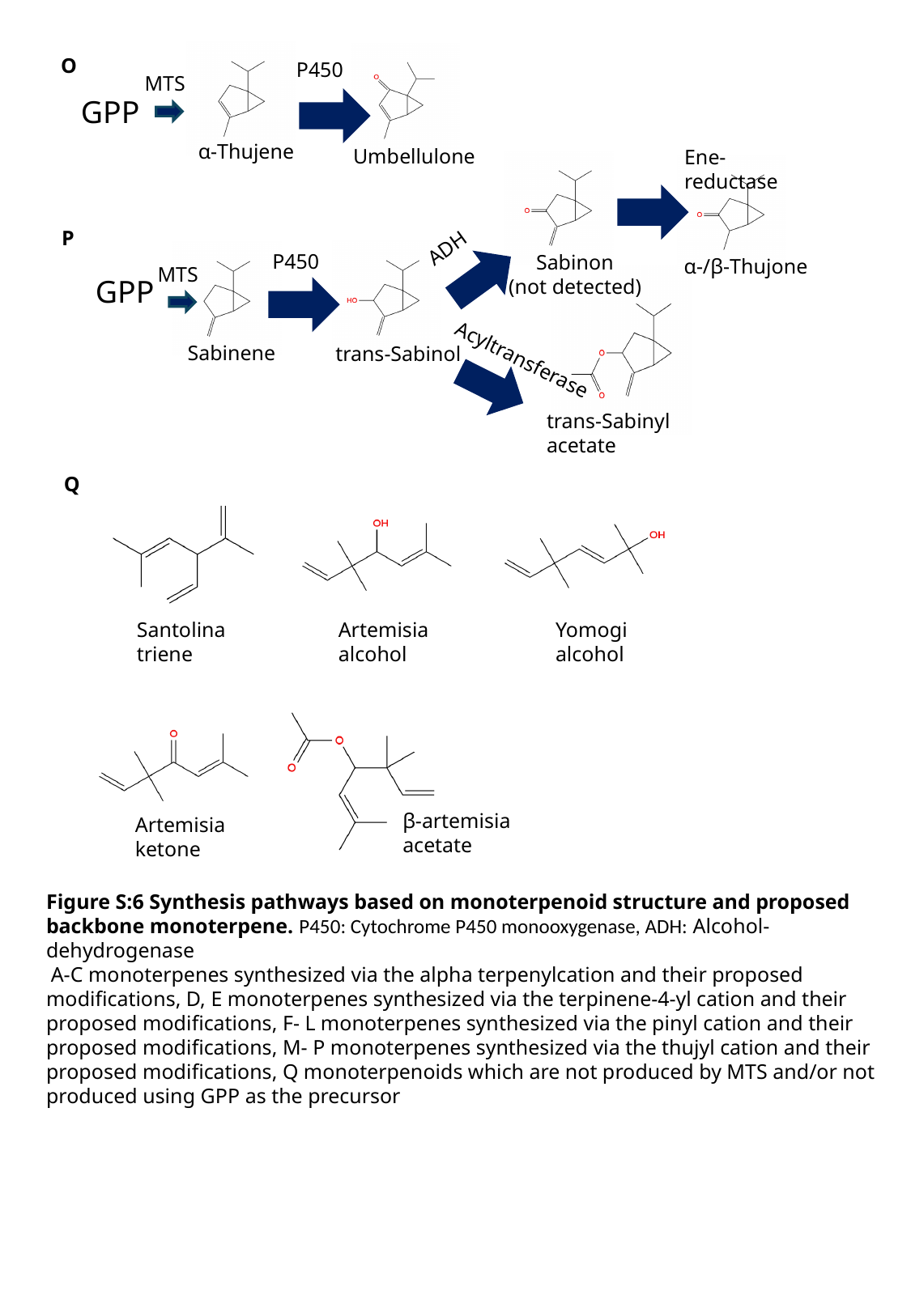

α-Thujene
Umbellulone
O
P450
MTS
GPP
Ene-reductase
Sabinon
(not detected)
α-/β-Thujone
ADH
P
trans-Sabinol
Sabinene
P450
MTS
trans-Sabinyl acetate
GPP
Acyltransferase
Q
Yomogi alcohol
Santolina triene
Artemisia alcohol
β-artemisia acetate
Artemisia ketone
Figure S:6 Synthesis pathways based on monoterpenoid structure and proposed backbone monoterpene. P450: Cytochrome P450 monooxygenase, ADH: Alcohol-dehydrogenase
 A-C monoterpenes synthesized via the alpha terpenylcation and their proposed modifications, D, E monoterpenes synthesized via the terpinene-4-yl cation and their proposed modifications, F- L monoterpenes synthesized via the pinyl cation and their proposed modifications, M- P monoterpenes synthesized via the thujyl cation and their proposed modifications, Q monoterpenoids which are not produced by MTS and/or not produced using GPP as the precursor

### Slide 10
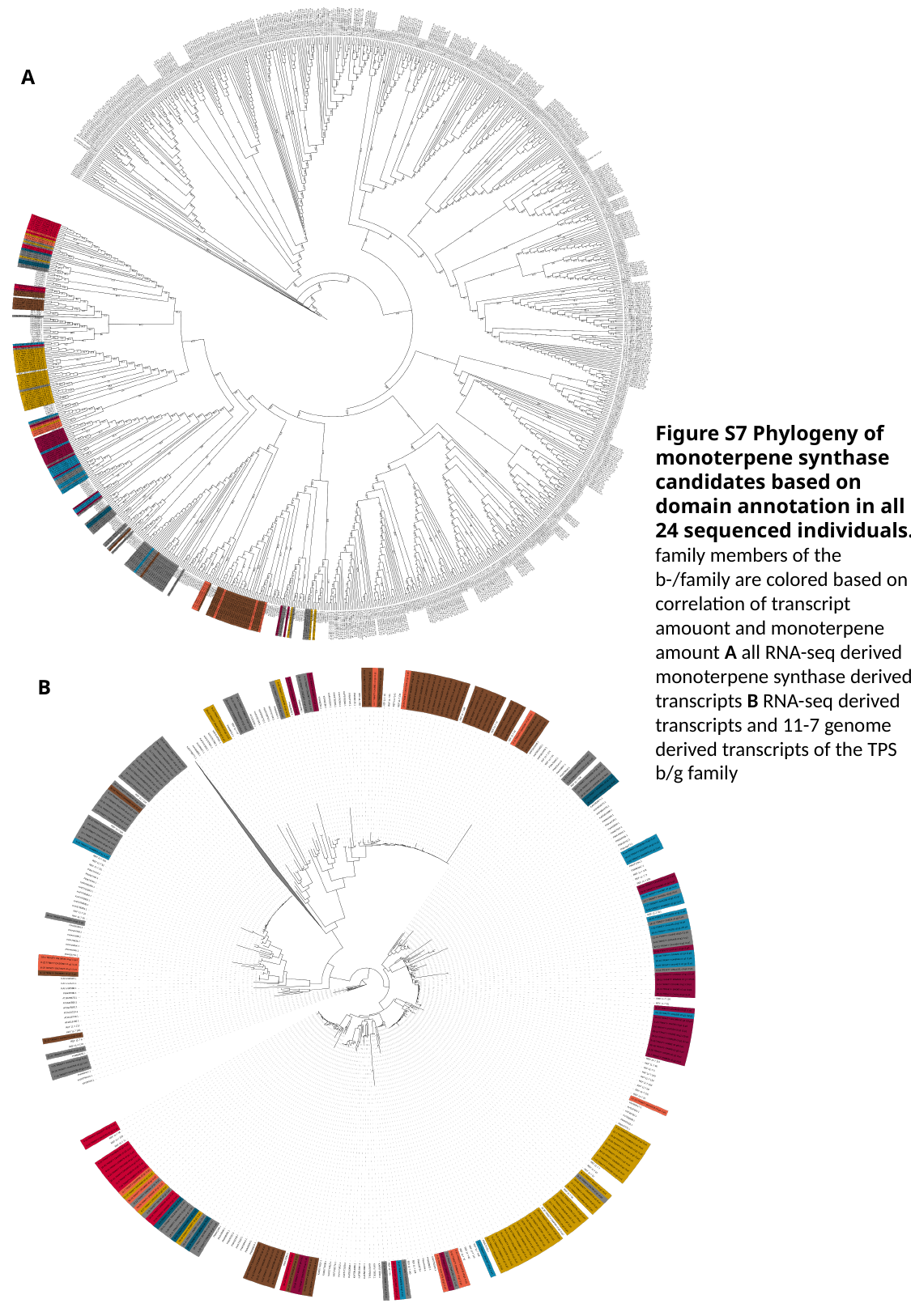

A
Figure S7 Phylogeny of monoterpene synthase candidates based on domain annotation in all 24 sequenced individuals. family members of the b-/family are colored based on correlation of transcript amouont and monoterpene amount A all RNA-seq derived monoterpene synthase derived transcripts B RNA-seq derived transcripts and 11-7 genome derived transcripts of the TPS b/g family
B

### Slide 11
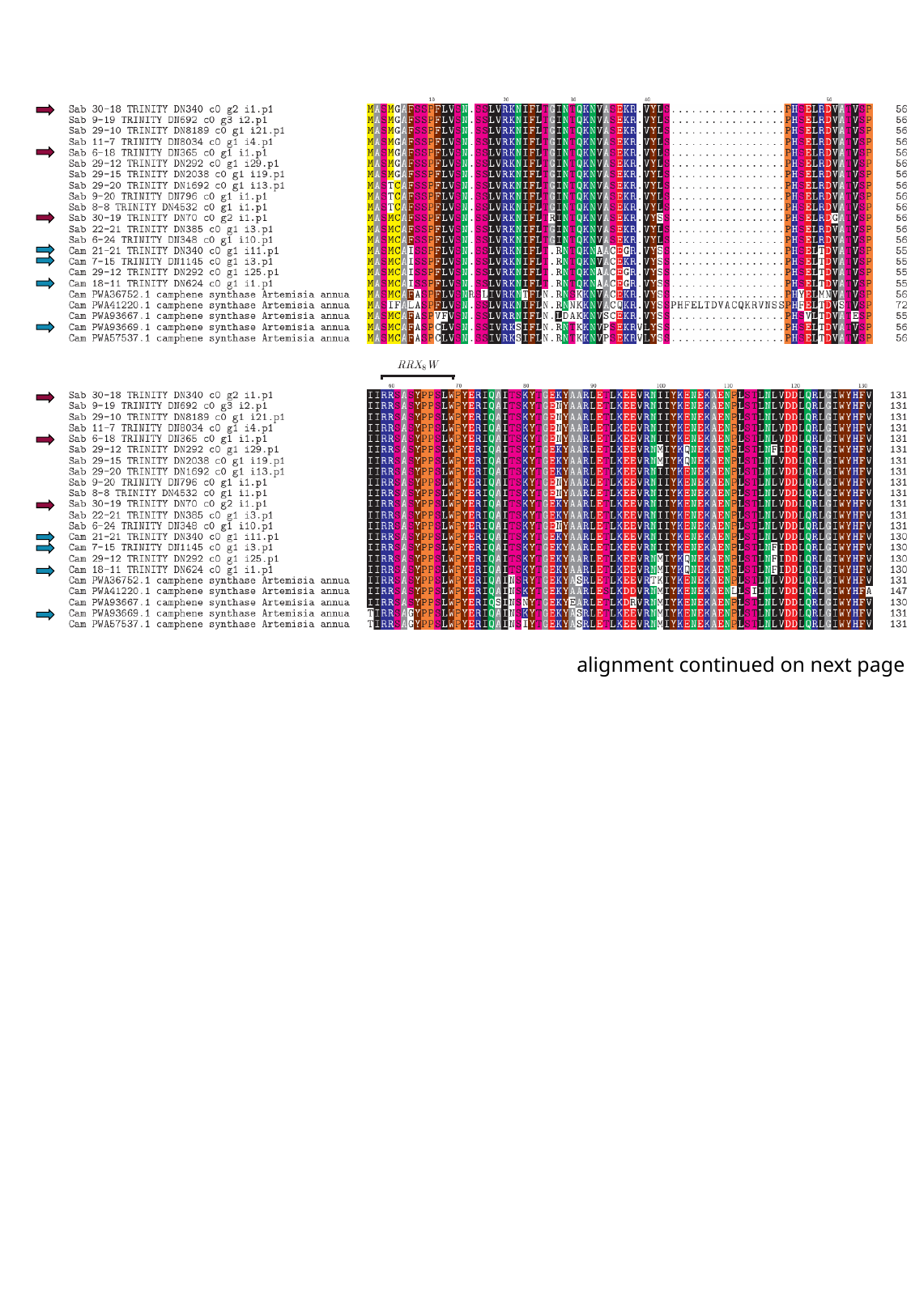

alignment continued on next page

### Slide 12
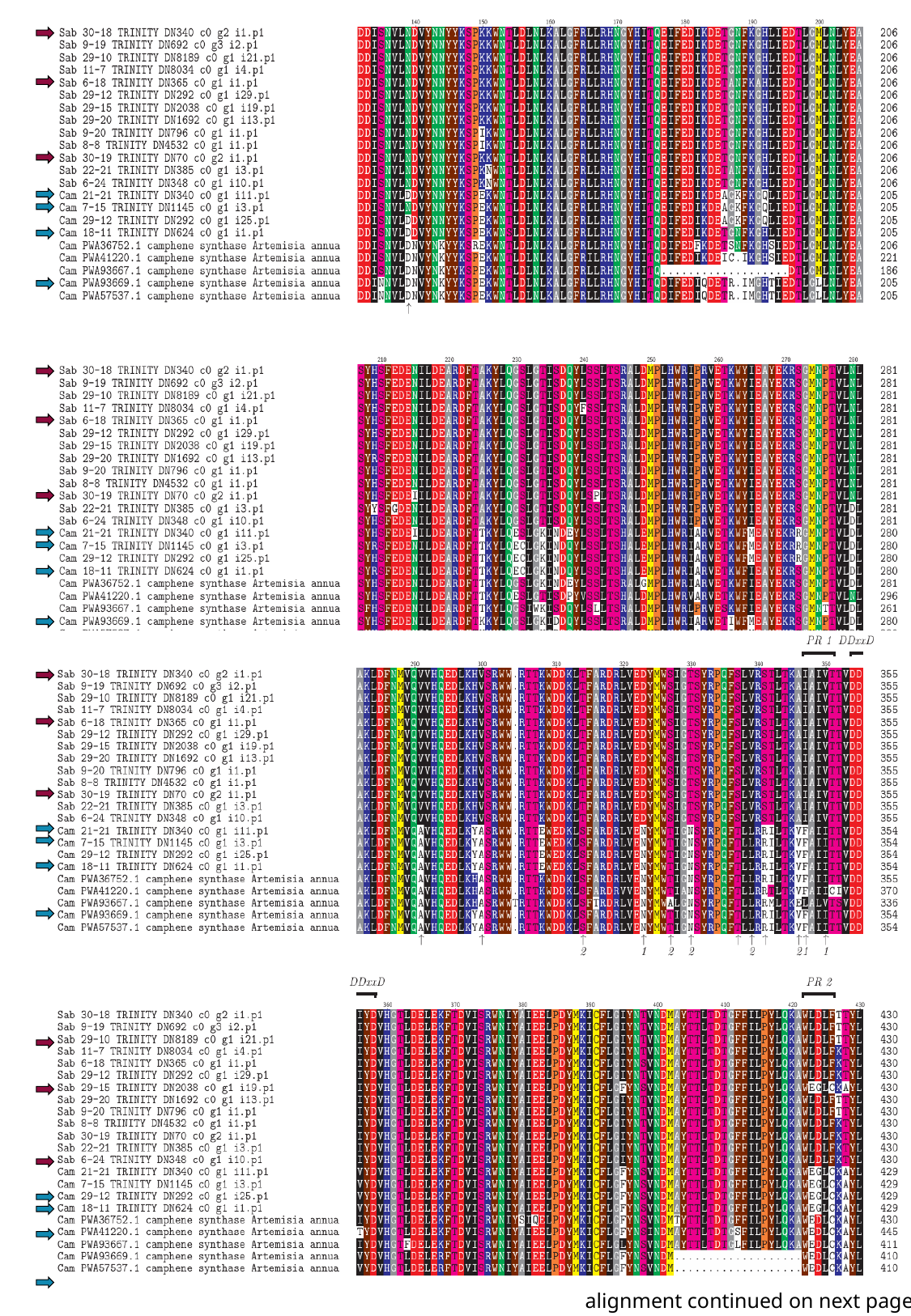

alignment continued on next page

### Slide 13
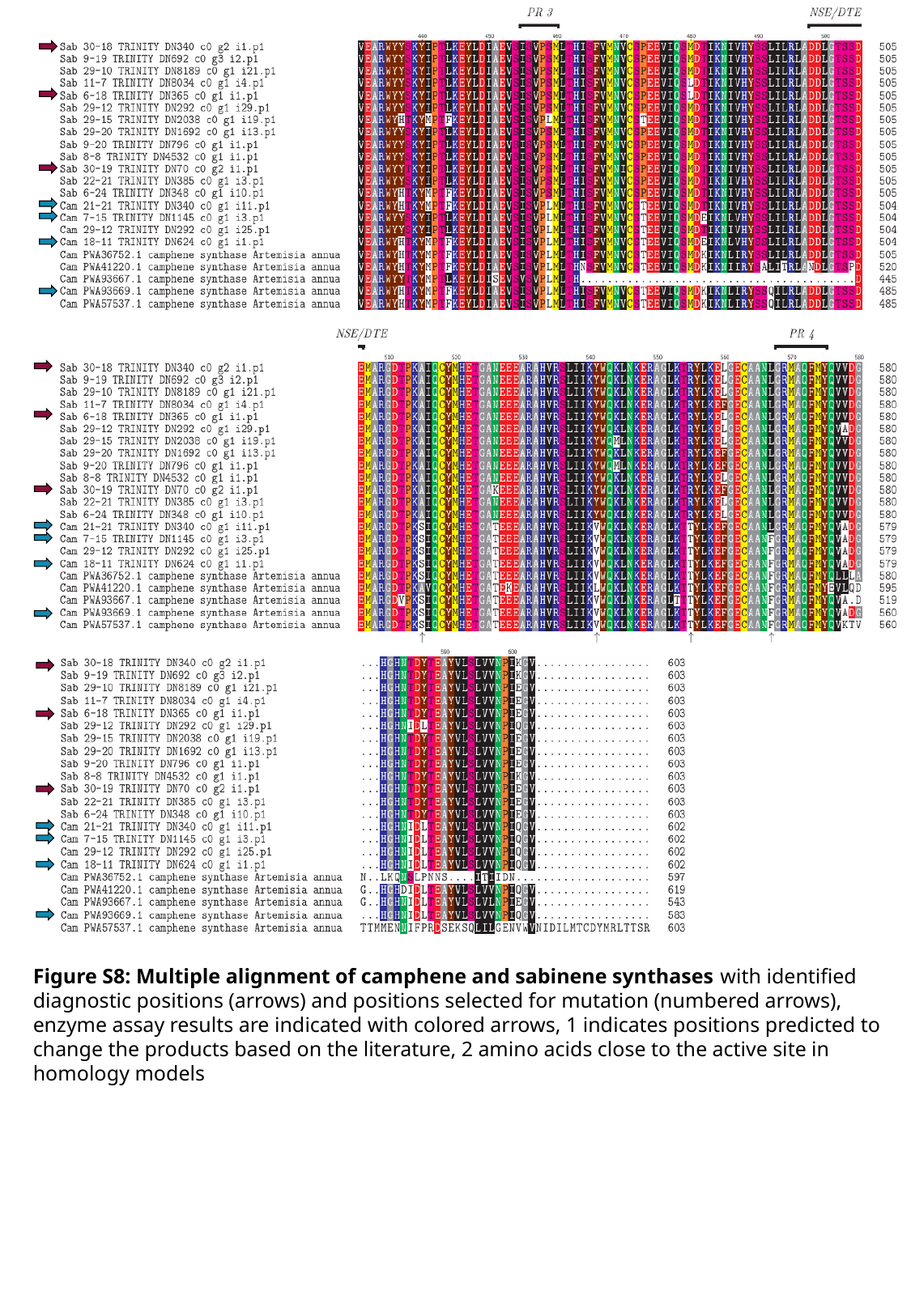

Figure S8: Multiple alignment of camphene and sabinene synthases with identified diagnostic positions (arrows) and positions selected for mutation (numbered arrows), enzyme assay results are indicated with colored arrows, 1 indicates positions predicted to change the products based on the literature, 2 amino acids close to the active site in homology models

### Slide 14
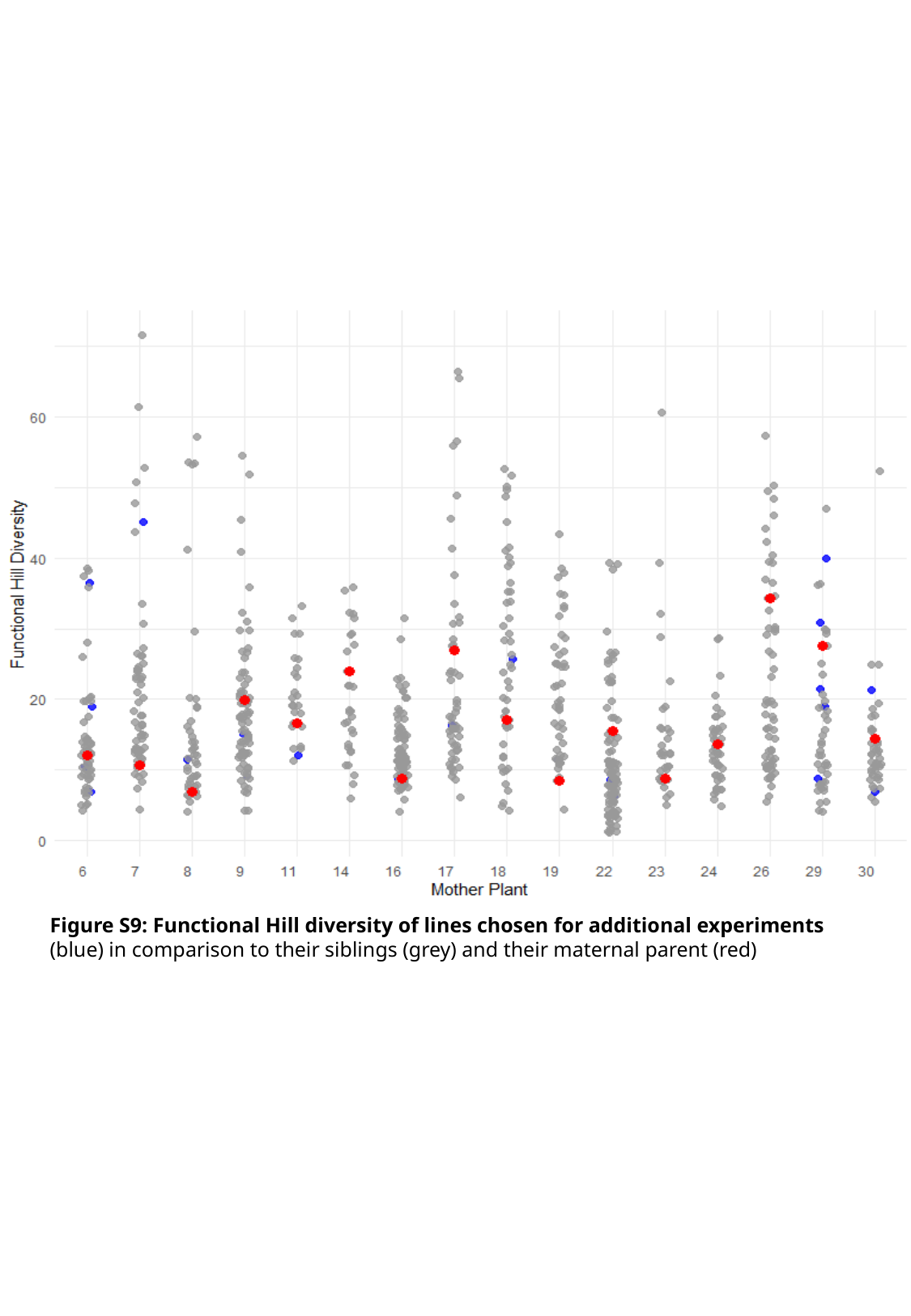

Figure S9: Functional Hill diversity of lines chosen for additional experiments (blue) in comparison to their siblings (grey) and their maternal parent (red)

### Slide 15
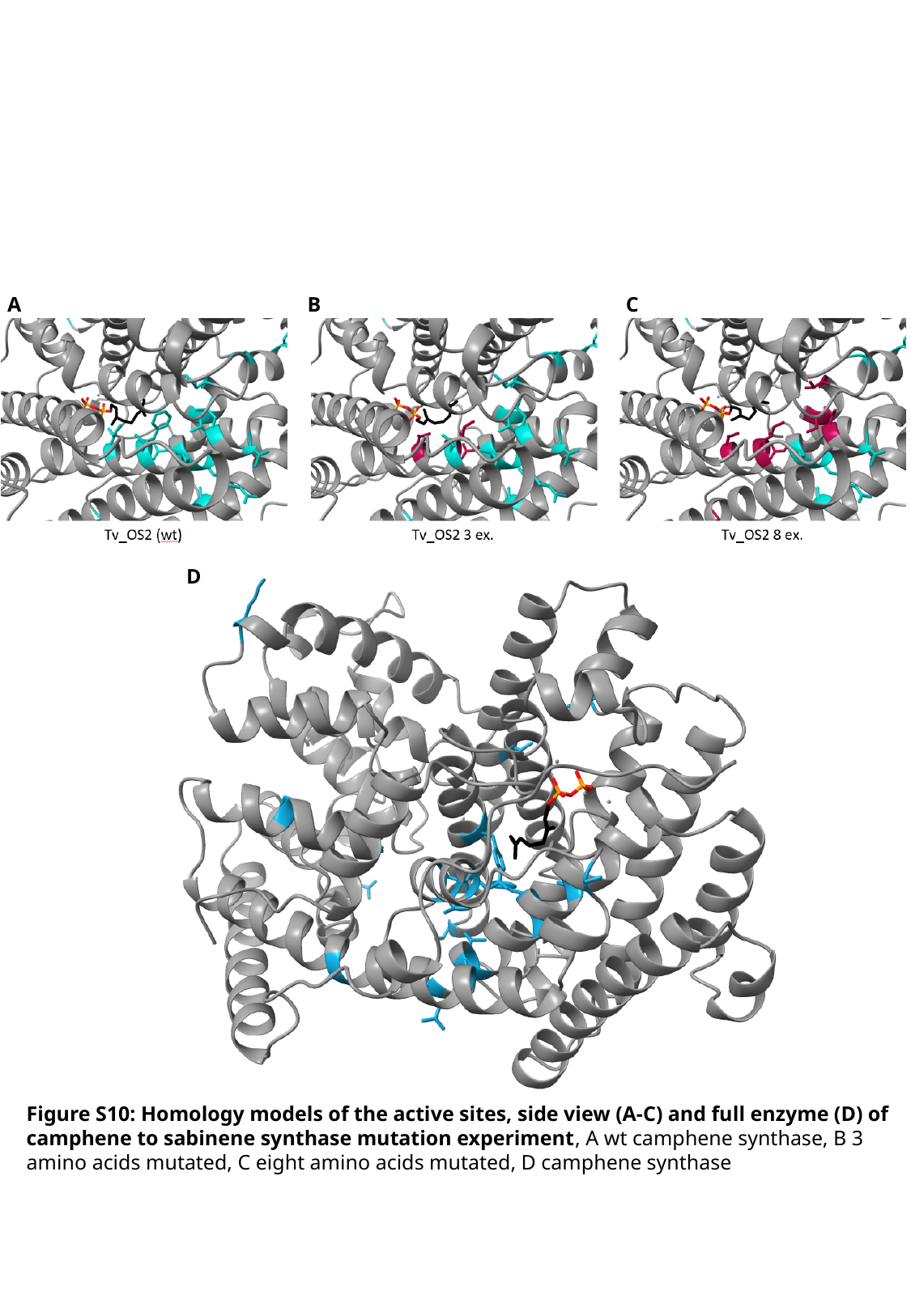

A
B
C
D
Figure S10: Homology models of the active sites, side view (A-C) and full enzyme (D) of camphene to sabinene synthase mutation experiment, A wt camphene synthase, B 3 amino acids mutated, C eight amino acids mutated, D camphene synthase

### Slide 16
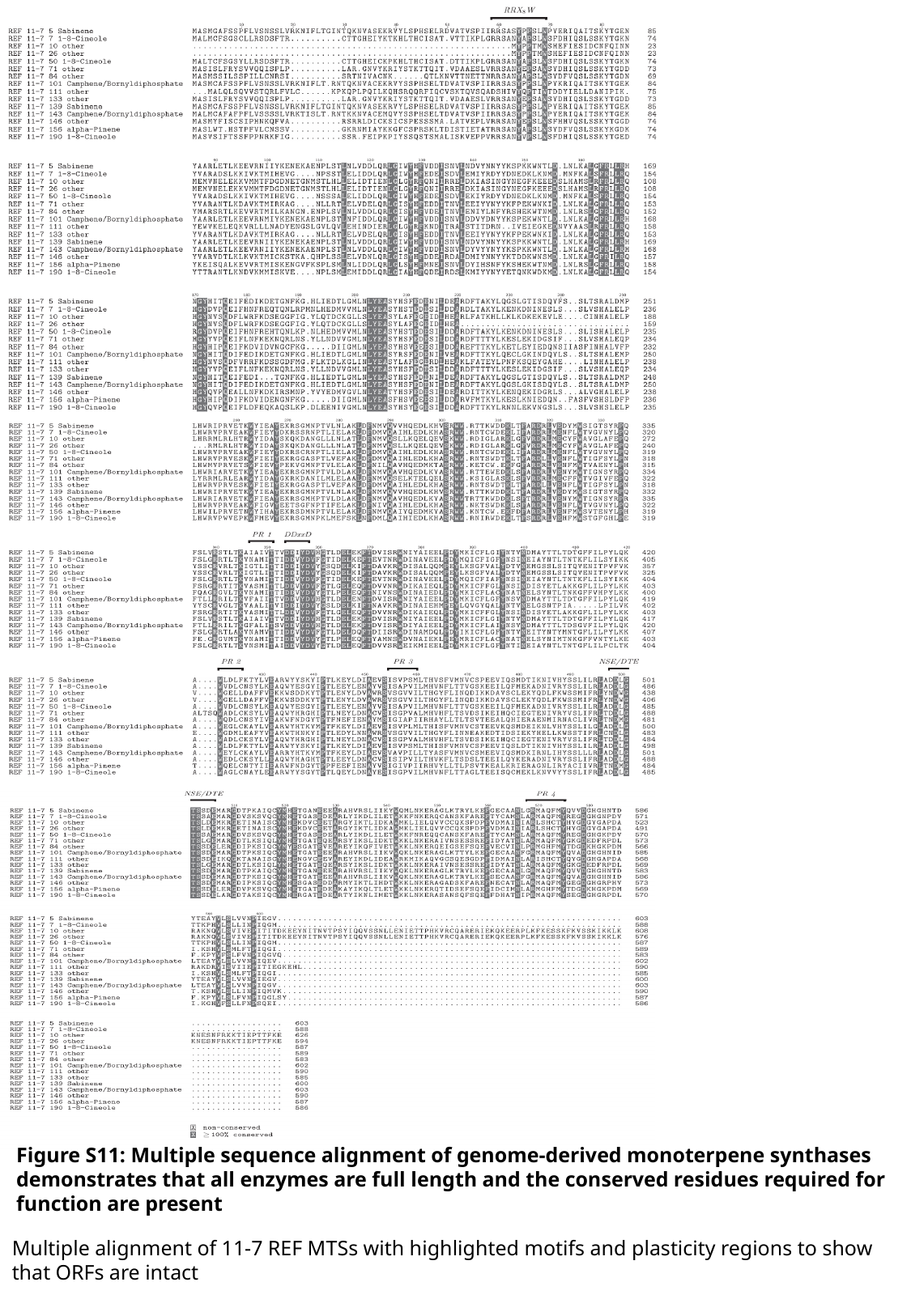

Figure S11: Multiple sequence alignment of genome-derived monoterpene synthases demonstrates that all enzymes are full length and the conserved residues required for function are present
Multiple alignment of 11-7 REF MTSs with highlighted motifs and plasticity regions to show that ORFs are intact

### Slide 17
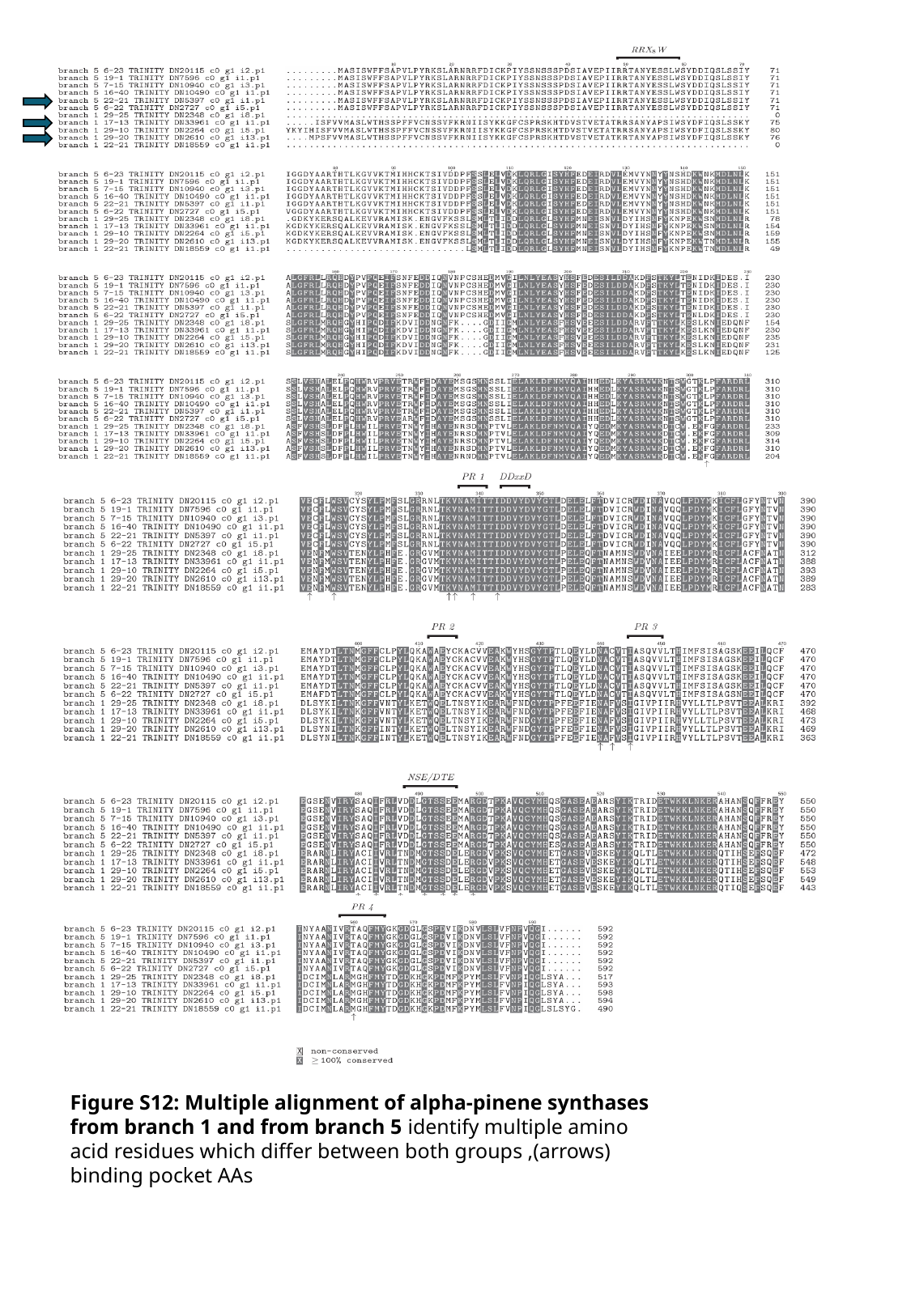

Figure S12: Multiple alignment of alpha-pinene synthases from branch 1 and from branch 5 identify multiple amino acid residues which differ between both groups ,(arrows) binding pocket AAs

### Slide 18
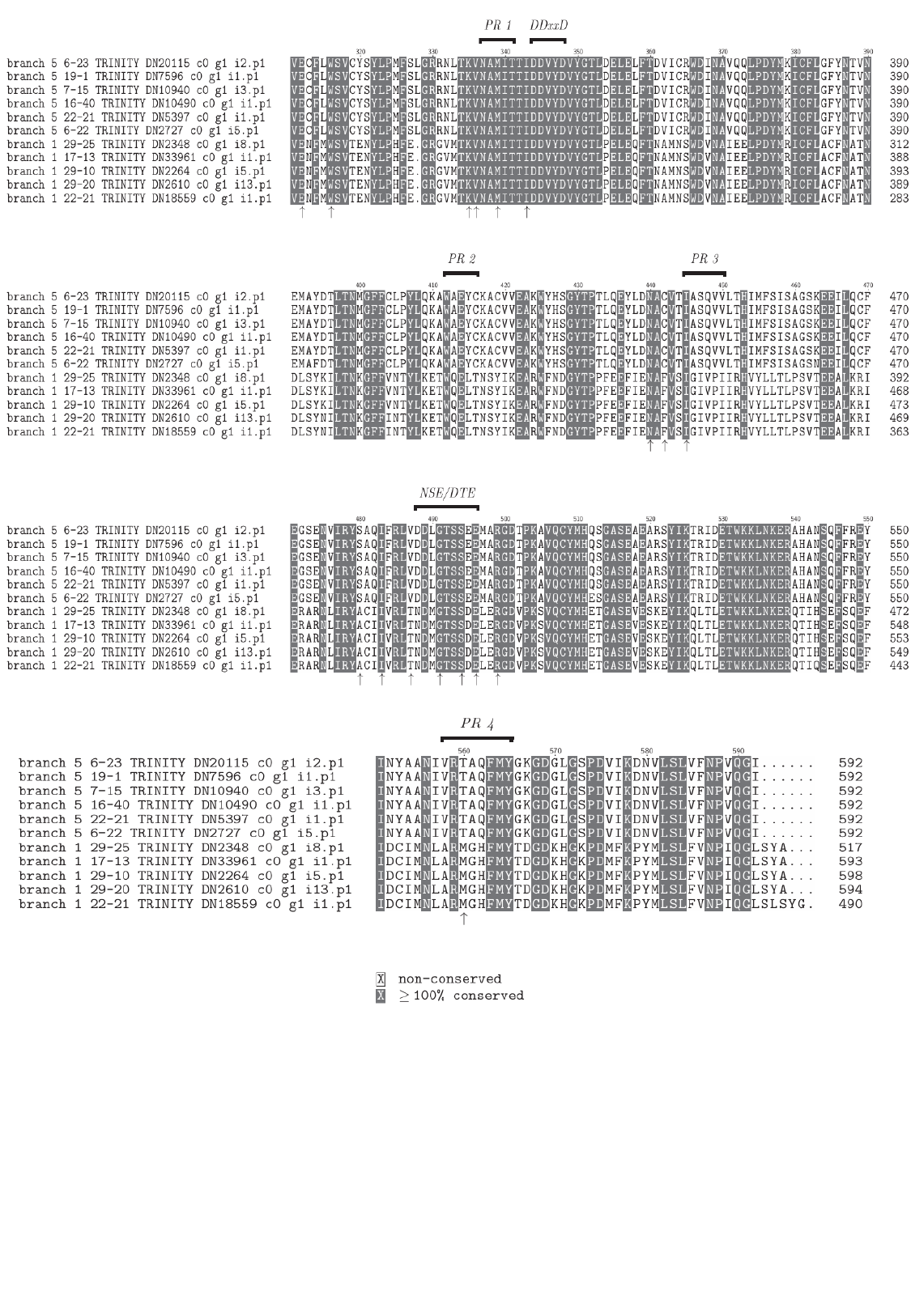
